# Type II collagen-positive progenitors are major stem cells to control skeleton development and vascular formation

**DOI:** 10.1101/2020.09.06.284588

**Authors:** Xinhua Li, Shuting Yang, Dian Jing, Lin Qin, Hu Zhao, Shuying Yang

**Author notes:** Corresponding author: Dr. Shuying Yang, Department of Basic and Translational Science, School of Dental Medicine, University of Pennsylvania, 240 South 40th Street, Levy 437, Philadelphia, PA 19104-6030.

## Abstract

Previous studies have revealed that type II collagen positive (Col2+) cells represent a kind of skeleton stem cells (SSC) and their descendants contribute to chondrocytes, osteoblasts, Cxcl12 (chemokine (C-X-C motif) ligand 12)-abundant stromal cells and bone marrow stromal/mesenchymal progenitor cells in postnatal life. To further elucidate the function of Col2+ progenitors, we generated mice with ablation of either embryonic or postnatal Col2+ cells. Embryonic ablation of Col2+ progenitors caused the mouse die at newborn with the absence of all skeleton except partial craniofacial bone, as well as multiple organ development defects and blood vessel loss. Postnatal ablation of Col2+ cells causes mouse growth retardation and collagenopathy phenotype. By examining Col2+ cells ablated mice, we found that, besides contributing to long bone and vertebral bone development, Col2+ cells are also involved in calvaria bone development. Meanwhile, Col2+ cells are the major cells to contribute all skeletal development including spine, rib and long bones. Moreover, our functional study provide evidence that intramembranous ossification is involved in craniofacial bone formation and long bone development, but not participates in spine development.

By performing lineage tracing experiments in embryonic or postnatal mice, we discovered that the presence of Col2+ progenitors not only within the bone marrow and growth plate (GP) but also within articular cartilage. Moreover, the number and differentiation ability of Col2+ progenitors were decreased with age in long bone and knee. Furthermore, fate-mapping studies revealed that Col2+ progenitors also contributed to CD31+ blood vessel endothelial development in calvariae bone, long bone and many organs. Interestingly, we found only 25.4% CD31+ blood vessel endothelial in long bone but almost all the CD31+ blood vessel endothelial in calvariae bone are differentiated from Col2+ cells. Consistently, postnatal Col2+ cells differentiated to both chondrocytes and CD31+ blood vessel endothelial cells during bone fracture healing. Therefore, this study reveals that Col2+ progenitors are the major source of endochondral ossification, and they also contribute to vascular development in multiple organs and fracture repair.

## Introduction

Multipotent mesenchymal stromal cells (MSCs) have been defined in culture as nonhematopoietic, plastic-adherent and colony-forming cells, which can differentiate into osteogenic, chondrogenic, and adipogenic progeny^1,2^. Recent studies have provided important insights into skeletal progenitor cells *in vivo* by using lineage tracing techniques^3^. Leptin receptor positive (Lepr+)^4,5^, chemokine (C-X-C motif) ligand 12 positive (Cxcl12+)^6^, Gli1+^7^, Prx+^8^, Nestin+^9^, type II collagen-positive (Col2+)^10,11^, Sox9+^11^, Aggrecan positive (Acan+)^11^ and Ctsk+^12^ cells have been reported to label MSCs and play important roles during bone development and homeostasis. Among these cells, the promoter/enhancer activities of Col2 were reported to label one of the widest cell populations *in vivo,* encompassing early mesenchymal progenitors that continue to become stromal cells, chondrocytes, osteoblasts and adipocytes^10,11,13^. Although Col2+ cells were regarded as early mesenchymal progenitors, questions remain regarding the function of Col2+ cells *in vivo*, how Col2+ progenitors change with age and what cell lineages Col2+ cells can contribute to.

Endochondral and intramembranous ossification are two processes for bone development in humans and mice. The main difference between endochondral ossification and intramembranous ossification is that endochondral ossification forms bone through a cartilage intermediate, while intramembranous ossification directly forms bone on the mesenchyme. It has been well known that most bones in the body are formed through endochondral ossification, and only the bones in the skull and clavicles ossify intramembranously^14,15^. However, thus far, there is no marker to identify endochondral ossification or intramembranous ossification. Recently, Debnath et al.^12^ discovered that Ctsk+ periosteal stem cells mediate intramembranous bone formation during bone development, but they still acquire endochondral bone formation capacity in response to injury. During the development stage, whether intramembranous ossification participates in long bone development and whether endochondral ossification participates in flat bone development are still unclear.

Here, to deeply understand the function of Col2+ cells, we generated genetic lineage tracing mouse models with tdTomato transgenic line and the mice with Col2+ cell ablation via diphtheria toxin (DTA) transgenic line. By analyzing these models in the embryonic and postnatal stages with age and determining the function of primary Col2+ cells in vitro, we provided the first evidence that Col2+ progenitors are the major source of endochondral ossification, and they also contribute to vascular development in multiple organs and fracture repair.

## Results

### Ablation of Col2+ cells causes mouse lethal at newborn with the absence of most of bone and cartilage

To generate the mice with the ablation of Col2+ embryonic progenitors, mice bearing a DTA transgene downstream of a floxed stop condon (DTA^fl/fl^) were crossed with mice expressing Col2-cre (mutant mice). Cre- littermates served as the control (wild type mice, WT mice). At embryonic day 17.5 (E17.5), Col2-cre; DTA^fl/−^ embryos can survival and follow Mendelian law (19/80). Interestingly, we found that more than one-quarter of the newborn (16/72) Col2-cre; DTA^fl/−^ mice were alive in the initial minutes or few hours following delivery but soon died because of oxygen insufficiency or severe organ development defects (Figure 1A, Supplementary figure 1A). The gross appearance of the Col2-cre; DTA^fl/−^ newborn mice showed severe developmental defects: severely dwarfish with extremely short four limbs and tail, intact but white appearance of the skin, cleft palate and abnormal skull morphology (Supplementary figure 1 B, C). Surprisedly, the eyes did not develop in some Col2-cre; DTA^fl/−^ newborn mice (9/16), indicating that Col2+ cells may contribute to mouse eye development.

**Figure 1.**
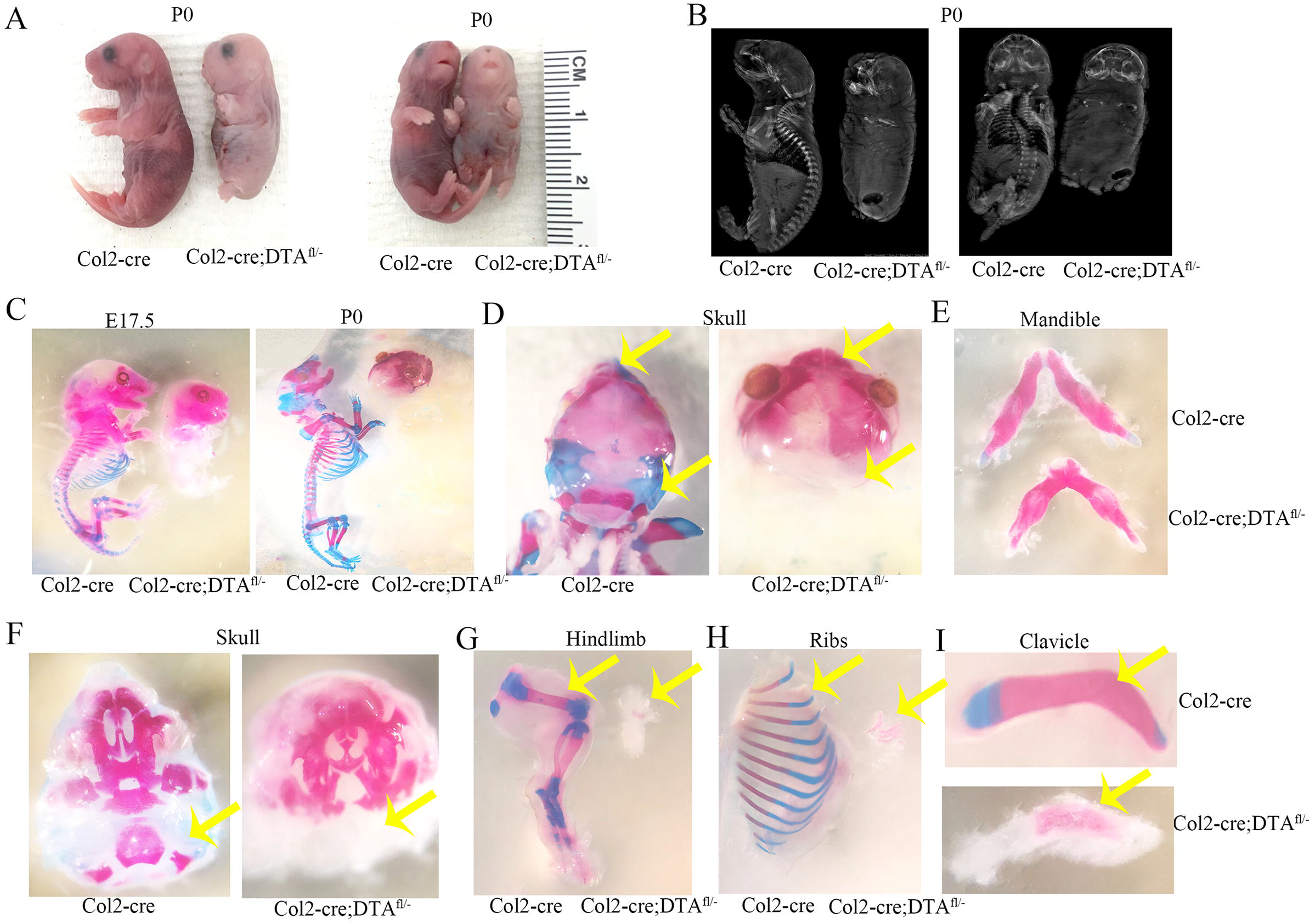
Ablation of Col2+ cells causes mouse lethal at newborn with the absence of endochondral bone and cartilage. (A)Gross appearance of Col2-cre, and Col2-cre;DTA^fl/−^ mice at P0. Mutant mouse is small and has extremely short four limbs pattern. (B)X-ray of P0 wild type and mutant embryo. (C)The skeleton at E17.5 and newborns of Col2-cre and Col2-cre;DTA^fl/−^ mice. Embryos and newborns were double stained with Alizarin red/Alcian blue. (D) The Alizarin red/Alcian blue staining of skull in Col2-cre and Col2-cre;DTA^fl/−^ newborns. Yellow arrow, cartilage completely lose in Col2-cre;DTA^fl/−^ mice. (E) The Alizarin red/Alcian blue staining of the mandible of Col2-cre and Col2-cre;DTA^fl/−^ at newborns mice. (F)The interior view lacking a mandibles of skull in Col2-cre and Col2-cre;DTA^fl/−^ newborns mice. Yellow arrow, bone loss in Col2-cre;DTA^fl/−^ mice. (G)The hindlimb of Col2-cre and Col2-cre;DTA^fl/−^ newborns mice. Yellow arrow, bone loss in Col2-cre;DTA^fl/−^ mice. (H)The ribs of Col2-cre and Col2-cre;DTA^fl/−^ newborns mice. Yellow arrow, bone loss in Col2-cre;DTA^fl/−^ mice. (I)The clavicle of Col2-cre and Col2-cre;DTA^fl/−^ newborn mice. Yellow arrow, parts clavicle lose in Col2-cre;DTA^fl/−^ mice. n=6 mice per condition from 3 independent experiments.

Skeletal radiograph examination of Col2-cre; DTA^fl/−^ newborns revealed that only parts of the skull, mandible, clavicle bone, and tiny hindlimb and ribs were calcified, and no calcification was found in vertebrate and other bones (Figure 1B). Consistent with the X-ray results, Alizarin red/Alcian blue staining of mutant embryos at E17.5 showed the absence of Alcian blue staining throughout the body and only limited Alizarin red staining in the craniofacial bone (Figure 1C). From E17.5 to birth in the mutant mice, no Alcian blue staining positive tissues were detected. However, calcification stained by Alizarin red was detected in parts of the frontal bone, parietal bone, temporal bone, maxillary bone, interparietal bone, and supraoccipital bone in the skull (Figure 1D, F), and almost all of the mandible (Figure 1E). Interestingly, very small pieces of bone in the hindlimb (Figure 1G), parts of the ribs (Figure 1H) and half of the clavicle bone are also Alizarin Red stained positive (Figure 1I).

### Ablation of Col2+ cells causes the absence of spine and disrupted all skeleton derived from endochondral ossification

To shed more light on the function of Col2+ progenitors in mice, we performed sagittal sectioning of the cre control (WT) and mutant newborns. In WT newborns, all the bone, cartilage, brain, spine cord and organs were well organized and clearly identified (Figure 2A). However, most bones, including the spine, were lost in mutant mice. Most interestingly, the brain and spine cord were well located in a relatively narrow skull cavity and “spinal canal”, respectively, despite severe bone loss in the skull and complete loss of vertebral bone and intervertebral disc in the mutant mice. However, the morphology of the spinal cord formed a spindle-like structure, indicating the spine or vertebral bone are needed to guide the normal spinal cord morphology.

**Figure 2.**
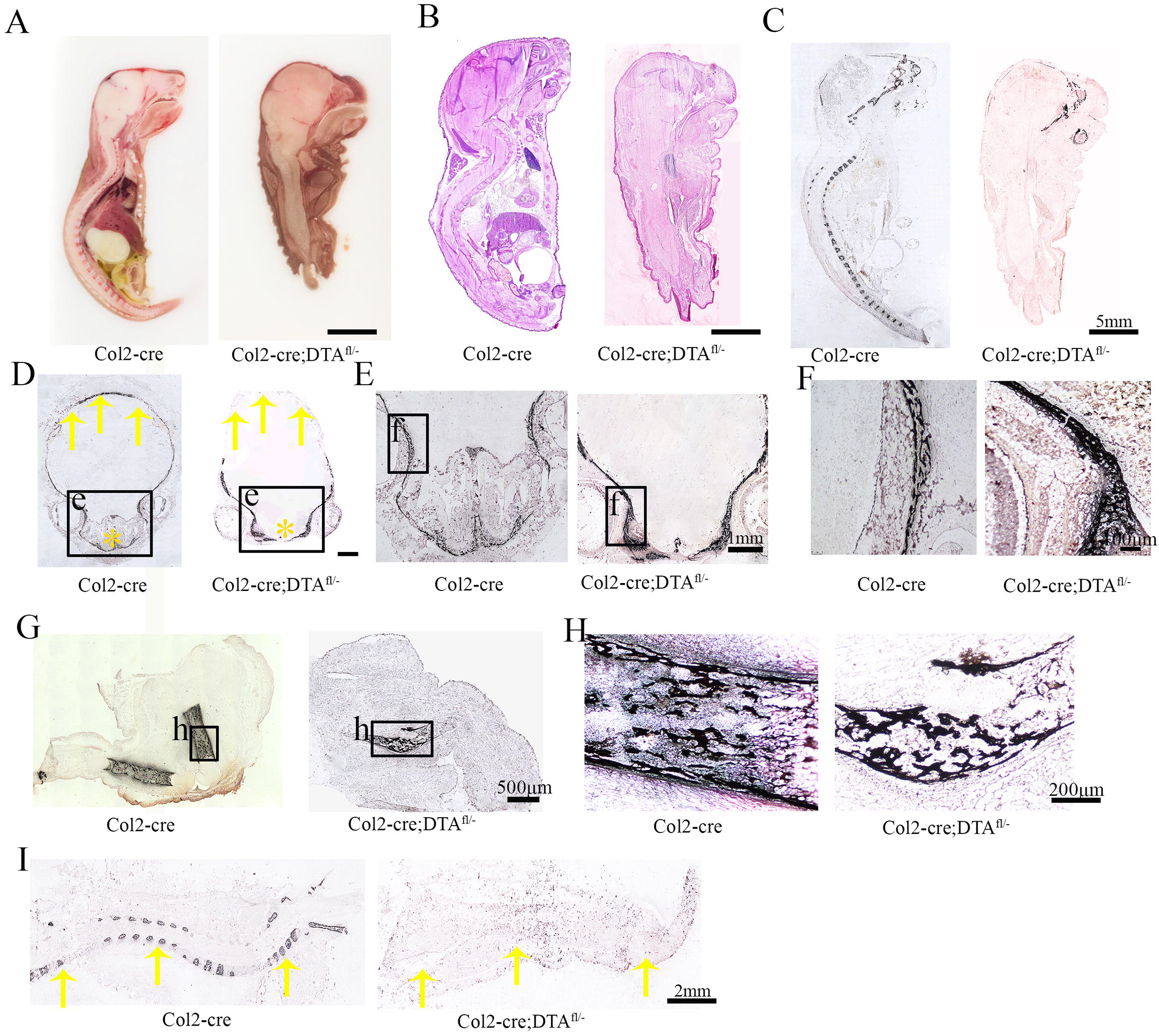
Ablation of Col2+ cells causes absence of spine and disrupts all endochondral skeletal development. Total view the middle sagittal section of Col2-cre and Col2-cre;DTA^fl/−^ mice at P0. H&E staining for the middle sagittal section of Col2-cre and Col2-cre;DTA^fl/−^ mice at P0. (C) Voncosa staining of the middle sagittal section of Col2-cre and Col2-cre;DTA^fl/−^ mice at P0. (D) Voncosa staining of the skull with transection. Yellow arrow, severe bone loss in back of skull of Col2-cre;DTA^fl/−^ mice. Yellow star, nasal cavity area in Col2-cre and Col2-cre;DTA^fl/−^ mice. (E) Voncosa staining of craniofacial bone show the loss of nasal cavity in mutant mouse. (F) Higher magnification of Voncosa staining showing similar sponge like bone structure in both wild type and mutant mice. (G) Voncosa staining of bone in hindlimb of wild type and mutant mice. (H) Higher magnification of Voncosa staining in hindlimb of wild type and mutant mice. Note that a small piece of well calcified sponge like bone structure can be detected of Col2+ cells ablation mice. (I) Voncosa staining in the middle sagittal section of spine in wild type and mutant mice. Yellow arrow, severe vertebral bone loss in Col2-cre;DTA^fl/−^ mice. n=6 mice per condition from 3 independent experiments.

To gain further insights into the changes in each tissue, histological sections of the skeletons from newborns were examined by staining with H&E (Figure 2B, Supplementary figure 1D, E, F) and Von Kossa’s staining (Figure 2C). Consistently, we found that ablation of Col2+ cells caused the absence of most parts of skeleton including complete loss of vertebral bone and intervertebral disc, all other body bones except parts of the skull and tiny hindlimb bone left in mutant mice. To determine the defects in finer detail, we evaluated sections of the skull, hindlimbs and spine. In WT newborns, the nasal cavity and intact skull were observed with well calcified bone. However, in mutant newborns, nasal cavity was absent, moreover, only the front of the calcified skull bone could be detected (Figure 2D, E, F), indicating that Col2+ cells play an important role in skull development. Most interestingly, a small piece of calcified sponge-like bone structure in the hindlimbs of Col2+ cell ablated mice could be detected (Figure 2G, H), indicating that besides endochondral ossification, intramembranous ossification may participate in long bone development. Von kossa staining result showed the clear vertebral bone mineralization in WT mice, which was completely absent in mutant mice, suggesting that Col2+ cells are the major progenitors contributing to spine bone development (Figure 2I). Most interesting, even though almost all bone and cartilage in mouse body were lost in mutant newborns, the toe and finger numbers and intact patterns are present and correct, suggesting that limb pattern development is independent of Col2+ cells in mice (Supplementary figure1B).

### Col2+ cells are present not only skeleton but also other important organs, and ablation of Co2+ cells impair those organ development

To test the efficacy of Cre-mediated DTA ablation, we generated Col2-cre; tdTomato and Col2-cre; DTA^fl/−^; tdTomato mice. Lineage tracing revealed that abundant tdTomato+ cells were detected in the spine, limbs, ribs, meninges, skull bone and some cells in internal organs, skin, fat tissues and brain of Col2-cre; tdTomato mice (Figure 3A). However, no tdTomato+ cells were detected in mutant mice compared with WT mice. Notably, we found that many cells in WT skull bone are tdTomato+, especially in the cartilaginous nasal capsule (Figure 3B), confirming that Col2+ cells contribute to skull bone and nasal capsule formation. Consistently, the cartilaginous nasal capsule was completely lost, and the skull was smaller in Col2-cre; DTA^fl/−^; tdTomato newborns than in control newborns. In the lower extremities and spine, most cells were labeled, although articular chondrocytes exhibited a weak signal (Figure 3C, D). In contrast, no tdTomato+ cells could be detected in the spine and lower extremities in Col2-cre; DTA^fl/−^; tdTomato newborns, confirming the effectiveness of the cell ablation.

**Figure 3.**
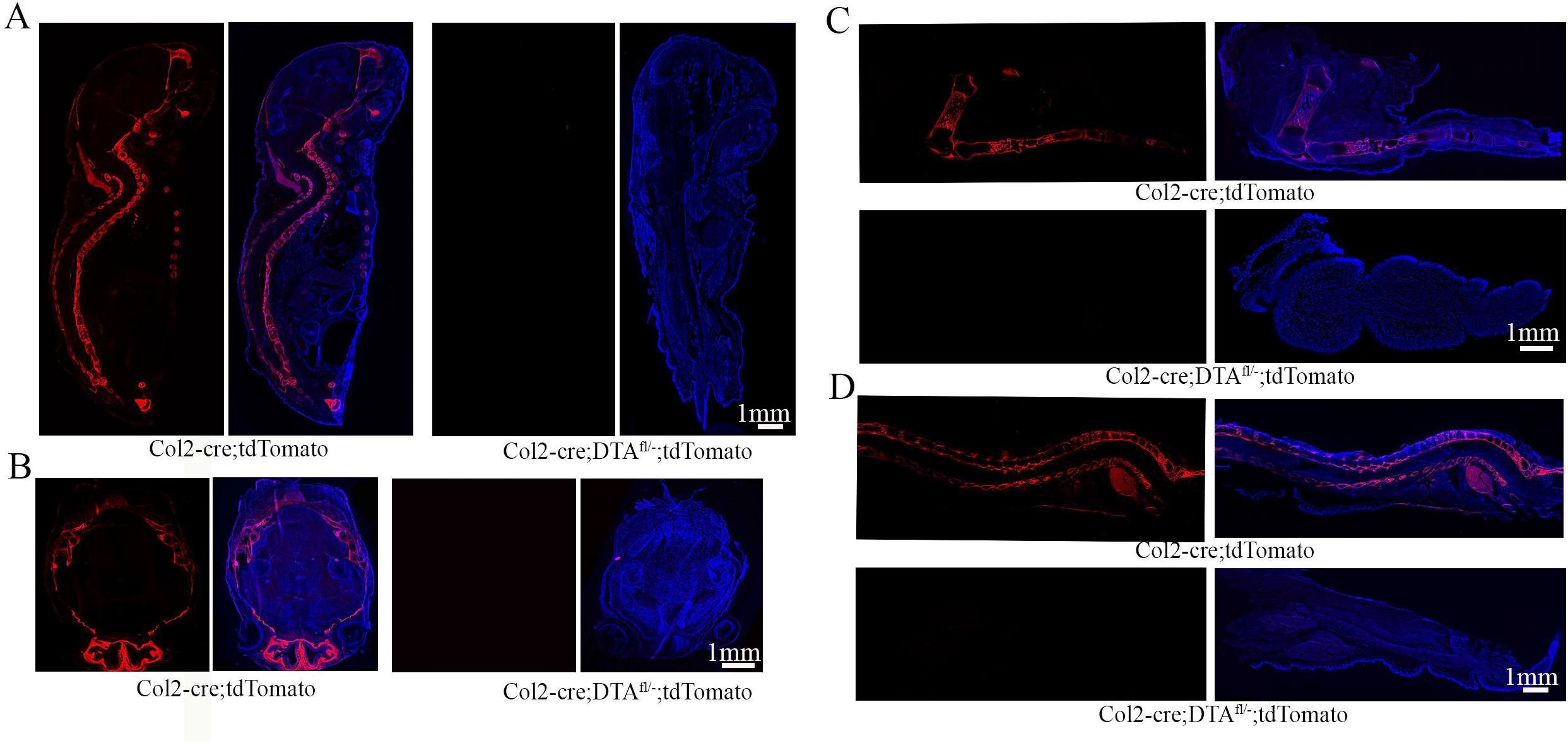
Spatial expression of Col2+ lineage progenitors in newborn and DTA are enough to remove the majority of Col2+ cells. The fluorescence images showing the pattern of Col2+ lineage progenitors in middle section of Col2-cre;tdTomato and Col2-cre;DTA^fl/−^;tdTomato newborn mouse. The fluorescence images showing the pattern of Col2+ lineage progenitors in transection of skull in Col2-cre;tdTomato and Col2-cre;DTA^fl/−^;tdTomato mouse. (C) The fluorescence images showing the pattern of Col2+ lineage progenitors in lower extremities of Col2-cre;tdTomato and Col2-cre;DTA^fl/−^;tdTomato newborn mouse. Note that completely loss of tdTomato+ cells in the long bone of Col2-cre;DTA^fl/−^;tdTomato compared to wild type newborn mouse. (D) The fluorescence images showing the pattern of Col2+ lineage progenitors in the middle sagittal section of spine of Col2-cre;tdTomato and Col2-cre;DTA^fl/−^;tdTomato new born mice. n=6 mice per condition from 3 independent experiments.

To gain more insight into the contribution of Col2+ cells to different organs, we detected the tdTomato+ fluorescence signal in the heart, lung, kidney, testes, liver, epitidimus, stomach and trachea in 4-week-old Col2-cre; tdTomato mice (Supplementary figure 2). In the heart and liver, less than 1% of the myocardial, epicardial or hepatic tissues were tdTomato+. In the trachea, tdTomato+ cells were located in cartilage tissues. In lung tissues, most tdTomato+ cells were detected in the region surrounding bronchiole epithelia, and few tdTomato+ cells were found in alveolar cells. Interestingly, approximately 3% of Col2+ cells were only located in the cortex renis in the kidney. In the spleen, many Col2+ cells can be detected, suggesting that Col2+ cells may play a role in spleen development and function. Very interestingly, some Col2+ cells formed loop-like structures in the testes and epitidimus, and many tdTomato+ cells were detected in the stomach epithelium.

Consistently, in Col2+ cell-ablated newborn mice, the spleen tissues were completely lost, and the heart, lung and stomach were smaller and abnormal compared with the control (Supplementary figure 3A-H). Histological results revealed that the heart ventricle was narrowed, and the myocardium was disorganized (Supplementary figure 3A, F). The alveoli were smaller, and most did not have an empty cavity compared with WT mice (Supplementary figure 3B, G). Abnormal kidney morphology can be found in Col2-cre; DTA^fl/−^ mice (Supplementary figure 3C, H). H&E staining showed that the kidney structure was dramatically disrupted without a clear boundary between the cortex renis and the medulla, and the cortex renis was severely defective in mutant mice. These findings demonstrate that Col2+ cells contribute other important organ development besides skeletal development.

### Postnatal deletion of Col2+ cells causes mouse growth retardation and a type II collagenopathy phenotype

To assess the contribution of postnatal Col2+ cells to mouse skeletal development, we genetically ablated these cells by inducing the expression of DTA postnatally. Specifically, we applied tamoxifen (TM) to either TM-inducible type II collagen Cre (Col2-creERT) or Col2-creERT; DTA^fl/fl^ mice at P3 of age and harvested the mice at 4 weeks of age. All Col2-creERT; DTA^fl/fl^ mice developed postnatal dwarfism with shorter limbs and body length (Figure 4A-E). Skeletal radiographs and whole-mount Alizarin red skeletal staining confirmed this observation and showed skeletal defects included epiphyseal dysplasia in the long bone and spine, flattened vertebral bodies, abnormalities of the capital femoral epiphyses and underdevelopment of the femoral head (Figure 4D, E). Representative micro-CT images showed epiphyseal dysplasia and dramatically decreased bone mass in the long bones of mutant mice (Figure 4 F, G). In particular, BV/TV, Tb.N, and Tb.Th were reduced to approximately 0.62-, 0.53-, and 0.5-fold, respectively, whereas Tb.Sp was increased 1.5-fold in Col2-creERT; DTA^fl/fl^ mice compared to those in Col2-creERT controls. Furthermore, Alizarin red/Alcian blue staining revealed a second ossification in the femur without any Alizarin red/Alcian blue positive staining in Col2-creERT; DTA^fl/fl^ mice (Figure 4H). Quantitative analysis showed the lengths of femur bones were significantly reduced from 1.22 cm in WT mice to 0.55 cm in the mutant group, and lengths of tibia bones were significantly reduced to 0.78 cm in the mutant group compare to 1.4 cm in WT mice (Figure 4I).

**Figure 4.**
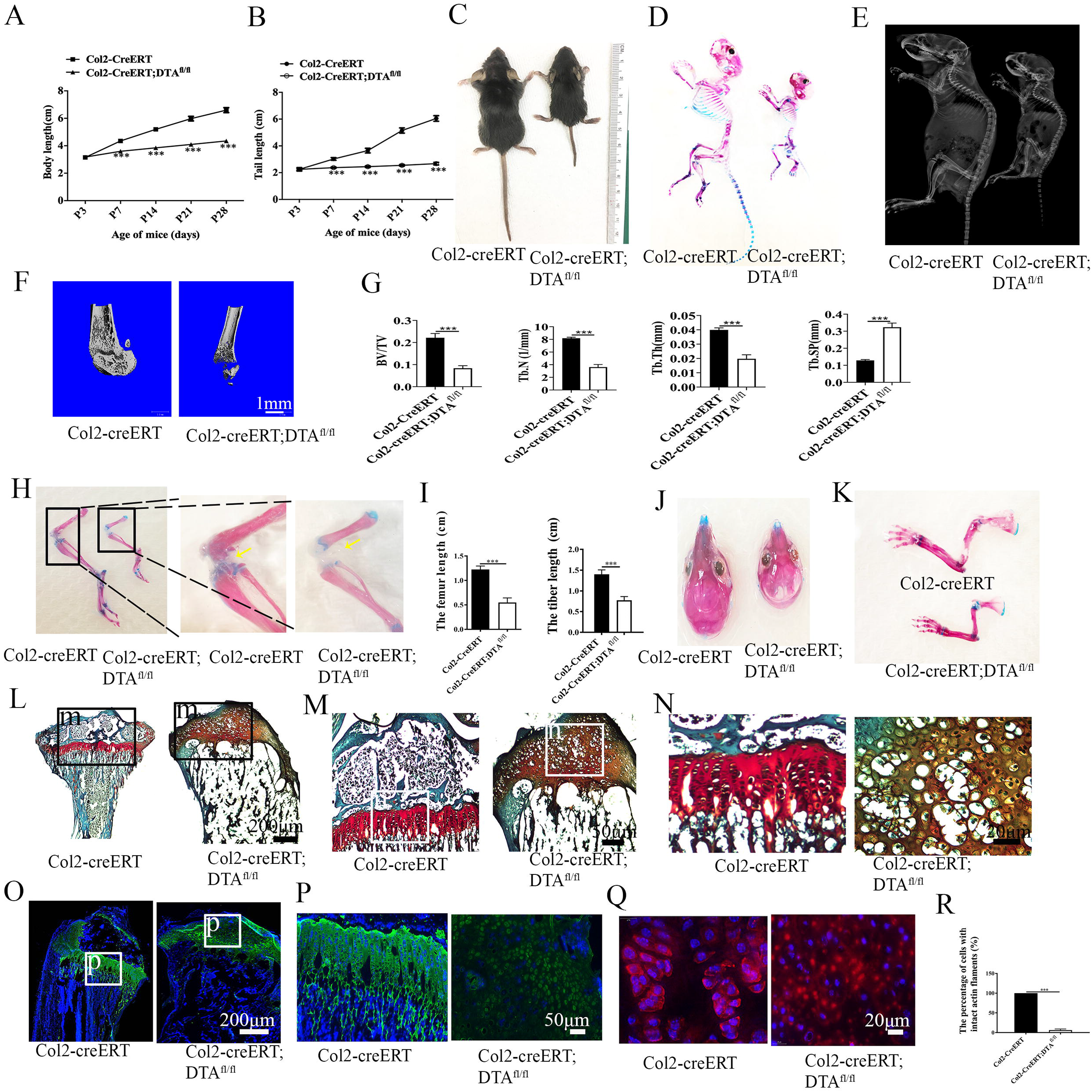
Postnatal deletion of Col2+ cells causes mouse growth retardation and a type II collagenopathy phenotype. (A, B) Quantity analysis of mice body length and tail length in Col2-creERT and Col2-creERT;DTA^fl/fl^ mice from P3 to 4-week-old. (n=6 mice per condition from 3 independent experiments). Data are mean ± s.d. (C) Macroscopic image of 4-week-old littermates. Mutant (right) mouse is small and dwarfism. (D) Total view of the Col2-creERT and Col2-creERT;DTA^fl/fl^ mice stained with double stained with Alizarin/Alcian blue at 4-week-old. (E) X-ray of 4-week-old Col2-creERT and Col2-creERT;DTA^fl/fl^ mice. (F)The representative picture of microCT 3D structure of femur head of 4-week-old Col2-creERT and Col2-creERT;DTA^fl/fl^ mice. (G)The quantity analysis of percentage of bone volume to total bone volume (BV/TV), trabecular thickness (Tb.Th), trabecular number (Tb.N), and trabecular spacing (Tb.Sp) in the femurs of 4-week-old Col2-creERT and Col2-creERT;DTA^fl/fl^ mice. (n=5 mice per group). Data are mean ± s.d. (H) Total view of low extremities in the Col2-creERT and Col2-creERT;DTA^fl/fl^ mice double stained with Alizarin red/Alcian blue at 4-week-old. Yellow arrow, delayed second ossification development in Col2-cre;DTA^fl/−^ mice. (I) The quantity analysis of mice femur length and tibia length of 4-week-old Col2-creERT and Col2-creERT;DTA^fl/fl^ mice. (n=6 mice per condition from 3 independent experiments). Data are mean ± s.d. (J)Total view of skull in the Col2-creERT and Col2-creERT;DTA^fl/fl^ mice stained with Alizarin red/Alcian blue at 4-week-old. (K) Total view of hand in the Col2-creERT and Col2-creERT;DTA^fl/fl^ mice stained with Alizarin red/Alcian blue at 4-week-old. (L) Safranin O/fast green staining of coronal sections of femur in 4-week-old Col2-creERT and Col2-creERT;DTA^fl/fl^ mice. (M) High magnification picture showed second ossification and cartilage in femur of 4-week-old Col2-creERT and Col2-creERT;DTA^fl/fl^ mice. (N) High magnification picture showing the chondrocyte morphology in 4-week-old Col2-creERT and Col2-creERT;DTA^fl/fl^ mice. (O) Immunofluorescence staining growth plate and articular cartilage for type 2 collagen in 4-week-old Col2-creERT and Col2-creERT;DTA^fl/fl^ mice. (P) High magnification picture showing type 2 collagen staining in 4-week-old Col2-creERT and Col2-creERT;DTA^fl/fl^ mice. (Q) Phalloidin staining show the cytoskeleton of cartilage in 4-week-old Col2-creERT and Col2-creERT;DTA^fl/fl^ mice. (R) Quantitative measurements of the percentage of cells with intact actin flaments to the total cells. (n=6 mice per condition from 3 independent experiments). Data are mean ± s.d. Statistical significance was determined by one-way ANOVA and Student’s t-test. *P < .05, **P < .01, ***P < .0001. NS = not statistically significant.

Thus, all these results indicate that postnatal Col2+ cells are important for skeletal development, and ablation of Col2+ cells in mice mimics the human type II collagenopathy phenotype^16,17^ (Figure 4 J, K) caused by the mutations of Col2a1 gene^17,18^, showing short-trunk dwarfism, and skeletal and vertebral deformities.

### Postnatal deletion of Col2+ cells disrupts endochondral ossification and cell alignment

To gain more insight into the skeletal change, we performed Safranin O/fast green staining of tibia sections from 4-week old Col2-creERT and Col2-creERT; DTA^fl/fl^ mice (TM injected at P3). The results further confirmed the significant decreased bone mass in the tibia and the absence of second ossification centers in Col2-creERT; DTA^fl/fl^ mice (Figure 4L). Higher magnification examination of secondary ossification center and GP in the tibia showed that the GP pattern dramatically disrupted, and disorganized hypertrophic chondrocytes occupied the epiphyses (Figure 4M, N).

Immunofluorescence staining examination for type II collagen showed that the collagen II matrix was enriched in articular cartilage and GP chondrocytes in 4-week-old WT mice (Figure 4O, P). However, very limited and weaker collagen II positive signaling was detected in epiphytic chondrocytes of Col2-creERT; DTA^fl/fl^ mice, indicating that Col2+ cells are essential for type II collagen production. Moreover, the result from F-actin staining showed that the cell alignment and pattern in GP chondrocytes were dramatically disrupted, and quantitative analysis showed the percentage of cells with intact actin filaments decreased from 100% in WT group to 9.2% in mutant mice. These results indicate that type II collagen matrix and Col2+ cells plays important roles in cell patterning and GP organization (Figure 4Q, R).

### Spatial distribution of embryonic and postnatal Col2+ cells in the mouse long bone and knee

To investigate the contribution of embryonic and postnatal Col2+ cells to the skeleton development, we traced the fate of Col2-expressing cells in Col2-cre; tdTomato, which shows Col2+ cells starting from embryonic stage, and in Col2-creERT; tdTomato mice which exbibit Col2+ cells after they were treated with TM at the postnatal selected times, by tracing for different time periods. In our system, Col2+ cells from both mouse strains and their descendants were permanently marked by the expression of red fluorescent protein tdTomato.

We first examined how embryonic Col2+ cells contribute to skeletal development in Col2-cre; tdTomato mice. In the long bone and knee joint, tdTomato+ cells were found in the articular cartilage (AC), GP, and bone surface, and osteocytes, as well as tendon and meniscus (Figure 5A). Interestingly, the tdTomato+ fluorescence signal is weaker at GP at P0 and P8 but stronger in the trabecular and cortical bone and meniscus. Notably, the tdTomato+ fluorescence is stronger in every compartment of the knee at P14. With aging, the tdTomato+ fluorescence signaling gradually decreased in the trabecular and cortical bone and kept strong in AC and GP.

**Figure 5.**
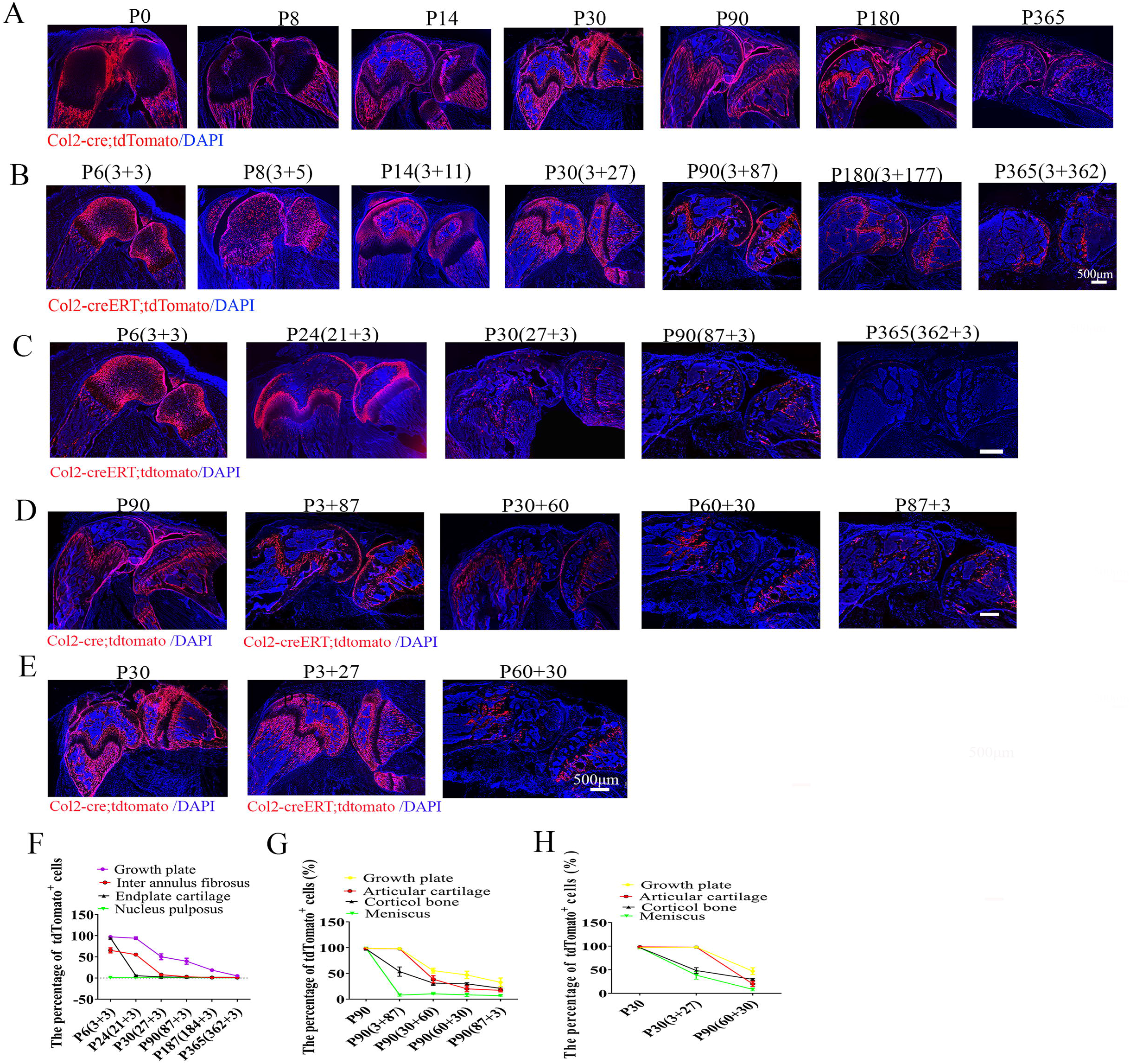
Spatial distribution of embryonic and postnatal Col2+ cells in the mouse long bone and knee. (A) Representative images from lineage tracing of embryonic Col2+ in long bone and knee joint at different time points (P0, P8, P14, P30, P90, P180 and P365). (B) Representative images from lineage tracing of postnatal Col2+ in long bone and knee joint at different time points (P6, P8, P14, P30, P90, P180 and P365). It was performed by injecting 75mg/kg tamoxifen into P3 mice. (C) Representative images from lineage tracing of postnatal Col2+ in long bone and knee joint which was activated at different time points (P3, P21, P27, P87, and P362). It was performed by injecting 75mg/kg tamoxifen into mice at indicated points. (D) Representative images from lineage tracing of Col2+ cells in long bone and knee joint at P90, which was activated at different time points (Embryonic stage, P3, P30, P60 and P87). (E) Representative images from lineage tracing of Col2+ cells in long bone and knee joint for 1 month while activated at indicated time points (Embryonic stage, P3, P60). It was performed by injecting 75mg/kg tamoxifen into Col2-creERT;tdTomato mice at different time points. (F) Quantitative measurements of the percentage of the tdTomato+ cells to the total cells in (C). (n=6 mice per condition from 3 independent experiments). Data are mean ± s.d. (G) Quantitative measurements of the percentage of tdTomato+ cells to the total cells in (D). (n=6 mice per condition from 3 independent experiments). Data are mean ± s.d. (H) Quantitative measurements of the percentage of tdTomato+ cells to the total cells in (E) (n=6 mice per condition from 3 independent experiments). Data are mean ± s.d.. It was measured at least 1000 cells each sample. Statistical significance was determined by one-way ANOVA and Student’s t-test. *P < .05, **P < .01, ***P < .0001. NS = not statistically significant.

To further examine how postnatal Col2+ cells contribute to long bone and knee development, Col2-creERT; tdTomato mice were intraperitoneally (i.p.) administered with TM at P3 and harvested at P6, P8, P14, P30, P90, P180, and P365. At P6 and P8, tdTomato+ (Col2+) cells were found predominantly in the AC, secondary ossification center, GP, meniscus (Figure 5B). Interestingly, at P14 and P30, the pattern of Col2 + cells in AC, GP and long bone was similar to that in Col2-cre; tdTomato, showing decreased tdTomato+ signaling in AC and GP, but increased signaling in the cortical bone and cancellous bone. At P90 and later time points, tdTomato+ signaling continuously existed and fluorescence intensity gradually decreased in GP and AC. Moreover, tdTomato+ cells significantly decreased and then disappeared in cortical bone and cancellous bone in both Col2-cre; tdTomato and Col2-creERT; tdTomato mice with age.

### Numbers and differentiation ability of Col2+ progenitors are decreased with age

To further explore the effect of age on the fate of Col2+ progenitors, Col2-creERT; tdTomato mice were i.p injected with TM at P3, P21, P27, P87, and P362, and sections were analyzed three days after each injection. As shown in Figure 5C, tdTomato+ cells were detected in articular chondrocytes, GP, cortical bone and the meniscus when TM was administered at P3 and detected 3 days later. The number of tdTomato+ cells in mice significantly decreased with age. Approximately 98.1% of GP chondrocytes were tdTomato+ in the mice with TM injection at P3, which decreased to 89.9% when TM was injected at P21 and then to 53%, 32.7%, 23.2%, and 23.2% when TM was injected at P30, P90, P187 and P365, respectively. Similarly, approximately 97.8% of articular chondrocytes were tdTomato+ in the mice that were injected with TM at P3, which decreased to 86.2%, 26.4%, 17.2%, 14.6%, and 9.1% when TM was injected at P21, P30, P90, P187 and P365, respectively (Figure 5C, F). In cortical bone, 26.1% of osteocytes were tdTomato+ in the mice with TM injection at P3, which decreased to 23.7%, 22.1%, 21.1%, 11.1%, and 5% when TM was injected at P21, P30, P90, P187 and P365, respectively. In the meniscus, 38.1% of cells were tdTomato+ in the mice that were injected with TM at P3, which decreased to 31.1%, 7.8%, and 6.9% when TM was injected at P21, P30, and P90, respectively, while only a few positive cells were detected when TM was injected at P187 or P365. The results demonstrated that the numbers of Col2+ progenitors are decreased with age.

To determine how age affects the number and differentiation ability of Col2+ cells, we compared the number of tdTomato+ cells in the knee at P90 when Col2+ cells were activated at the embryonic stage in Col2-cre; tdTomato mice or at P3, P30, P60, and P88 with TM injection in Col2-creERT; tdTomato mice (Figure 5D). The number of tdTomato+ cells gradually decreased from the embryonic stage, P3, P30, P60 to P88. Specifically, at P90, 98.5% of GP chondrocytes were tdTomato+ when Col2+ was activated at the embryonic stage, however, those cells decreased to 98.2%, 55.4%, 47.1%, and 37.2% in Col2-creERT; tdTomato mice when TM was injected at P3, P30, P60 and P87, respectively (Figure 5D, G). Similarly, approximately 98.4% of articular chondrocytes were tdTomato+ when Col2+ was activated at the embryonic stage, which decreased to 97.9%, 38.7%, 20.1%, and 17.2% when TM was injected at P3, P30, P60 and, and to 17.2% at P87, respectively. In cortical bone, 97.6% of cells were tdTomato+ when Col2+ was activated at the embryonic stage, which decreased to 53.5%, 31.2%, 29.8%, and 21.1% when TM was injected at P3, P30, P60 and P87, respectively. In the meniscus, 98.8% of cells were tdTomato+ when Col2+ was activated at the embryonic stage, which decreased to 8.1%, 10.4%, 8.4%, and 6.9% when TM was injected at P3, P30, P60 and P87, respectively. These results suggest that the numbers and differentiation ability of Col2+ cells decrease during the aging process.

To further confirm the differentiation ability of Col2+ progenitors decreased with age, we compared the tdTomato+ cells in the knee during 1 month of tracing when Col2+ cells were activated at the embryonic stage, P3 and P60 (Figure 5E). Approximately 98.4% of articular chondrocytes were tdTomato+ when Col2+ was activated at the embryonic stage, which decreased to 97.9% and 38.7% when TM was injected at P3 and P60, respectively (Figure 5E, H). Similarly, 98.5% of GP chondrocytes were tdTomato+ when Col2+ was activated at the embryonic stage, which decreased to 98.2% and 55.4% when TM was injected at P3 and P60, respectively. In cortical bone, 97.6% of cells were tdTomato+ when Col2+ was activated at the embryonic stage, which decreased to 53.5% and 31.2% when TM was injected at P3 and P60, respectively. In the meniscus, 98.8% of cells were tdTomato+ when Col2+ was activated at the embryonic stage, which decreased to 8.1% and 1.3% when TM was injected at P3 and P60, respectively. These results further confirmed that the differentiation ability of Col2+ progenitors decreased with age.

Besides these, by comparing the type II collagen expression between embryonic and postnatal Col2+ cells, we found that the expression pattern of type II collagen was similar to the pattern of Col2+ cells at P6 in Col2-creERT; tdTomato mice with TM injection at P3. However, type II collagen expression was completely different to the patterns of Col2+ cells at P30, P180 and P365 in Col2-cre; tdTomato mice or Col2-creERT; tdTomato mice with i.p. TM injection at P27, P177 or P362 (Supplementary figure 4A-D). These result thus further indicated that Col2+ cells gradually decreased with age.

### Col2+ progenitors contribute to blood vessel formation in multiple organs

Consistent with previous studies^11^, we also found that Col2+ cells (tdTomato+) were present in GP, AC and bone in 4-week-old Col2-cre; tdTomato mice (Figure 6A). Surprisedly, we found that some CD31+ cells in blood vessels of some organs or tissues were also labeled with tdTomato+ signaling, including long bone and skull bone (Figure 6B, C, D, E, F). In the long bone’s bone marrow of 4-week-old Col2-cre; tdTomato mice, approximately 25.4% of CD31+ cells were Col2+ (Figure 6 B, C), while almost all the CD31+ cells were Col2+ in skull bone. Additionally, we found almost all the CD31+ blood vessels cells in the brain, eyeball, heart, skin and skull bone were Col2+ (Figure 6D, Supplementary figure 5A-D), whereas, none CD31+ cells are Col2+ in kidney vasculature during development. Consistently, we found ablation of Col2+ cells led to dramatically decrease of CD31+ blood vessel endothelial cells in the skull bone and eyes of Col2-cre; DTA^fl/fl^; tdTomato mice (Supplementary figure 5E, F). Thus, we found for the first time that Col2+ progenitor cells can differentiate into CD31+ cells and blood vessel formation in multiple tissues and organs. The different contributions of Col2+ cells to blood vessels in long bone and skull bone revealed that the cell origin of blood vessels in long bone and skull bone may differ.

**Figure 6.**
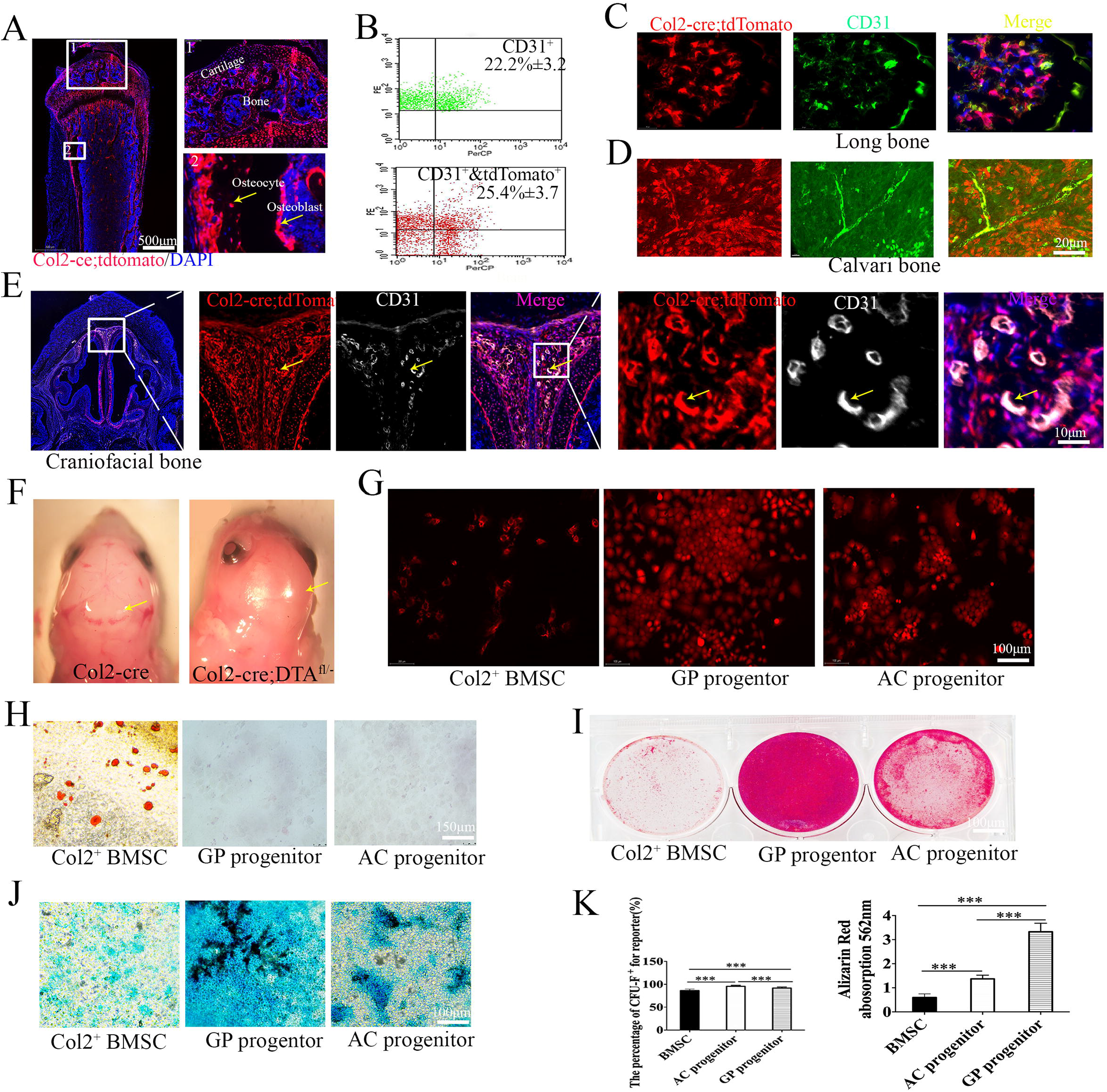
Col2+ progenitors have muti-lineage differentiation ability including CD31+ blood vessel. (A) Representative fluorescent images of distal femur showing cartilage, osteoblast and osteocyte are Col2+. Yellow arrow, osteoblast and osteocyte in 4 week-old Col2-cre;tdTomato mice. (B) Flow cytometry analysis was performed using dissociated bone marrow cells collected from 4-week-old Col2-cre; tdTomato mice. Representative dot plots showed some Col2+ bone marrow stromal cells are CD31+ (n=3 mice per condition from 3 independent experiments). (C) Representative fluorescent images from 4-week-old Col2-cre; tdTomato mice distal femur showing parts of CD31+ cells in long bone are Col2+. (D) Representative fluorescent images from 4-week-old Col2-cre; tdTomato mice calvaria bone showing almost all CD31+ cells are Col2+. (E) Representative fluorescent images from 4-week-old Col2-cre; tdTomato mice craniofacial bone showing almost all CD31+ cells are Col2+. Yellow arrow, the overlap staining of blood vessel. (F) The gross appearance of the head of Col2-cre and Col2-cre;DTA^fl/−^ newborn mice. Note that mutant mouse dramatically loss of blood vessel in skull. Yellow arrow, blood vessels in each group. (G) CFU-F assay in Col2+ MSC, GP progenitor and AC progenitor showed that every cell type can form CFU colons. (H) Osteogenic differentiation of Col2+ MSC, GP progenitor and articular cartilage (AC) progenitor. (I) Chondrogenesis differentiation of Col2+ MSC, GP progenitor and AC progenitor. (J) Adipogenesis differentiation of Col2+ MSC, GP progenitor and AC progenitor. (K) Quantitative measurements of CFU-F colonies and osteogenesis differentiation ability in (G and H). (n=3 mice per condition from 3 independent experiments). All data are reported as the mean ± s.d. Statistical significance was determined by one-way ANOVA and Student’s t-test. *P < .05, **P < .01, ***P < .0001. NS = not statistically significant.

### Col2+ cells in GP and AC display stem cell properties

Our results showed that Col2+ cells are located in AC, bone marrow cells (BMMs) and GP, and ablation of Col2+ cells completely disrupt skeletal development except craniofacial development. To determine whether Col2+ cells have stem cell properties, we first performed CFU-F activity assay. BMMs, GP chondrocytes and articular chondrocytes were isolated from 4-week-old Col2-cre; tdTomato mice, and then Col2+ cells were sorted out as shown in Figure 6G. Quantitation of Tomato+ CFU-F colonies revealed that Col2+ cells from BMMs, GP chondrocytes and articular chondrocytes can form CFU-F colonies (Figure 6G).

To further assess whether those Col2+ cells have multiple lineage differentiation capability, Col2+ cells from BMMs, GP and AC were respectively induced with osteogenic, chondrogenic and adipogenic media for indicated time. Interestingly, we found that Col2+ cells sorted from BMMs can differentiate into osteoblasts, chondrocytes and adipocytes, however, Col2+ cells from GP and AC can only differentiate into osteoblasts and chondrocytes, and cannot differentiate into adipocytes (Figure 6H-K), suggesting that Col2+ cells from BMMs are likely earlier stage of mesenchymal stem cells (MSCs), while a large population of Col2+ cells from GP and AC are later stages of MSCs. Most interestingly, quantitative analysis showed that Col2+ cells from GP have much stronger osteogenic and chondrogenic differentiation potential than Col2+ cells from articular chondrocytes and BMMs (Figure 6 H, L). It was reported that both blood vessel endothelial and blood cells are differentiated from hemangioblasts of mesodermal cells^14^. To identify whether Col2+ cells in bone marrow contribute to osteoclast formation, BMMs from Col2-cre; tdTomato mice were induced with RANKL/M-CSF. As shown in Supplementary Figure 6, tdTomato fluorescence was not present in osteoclasts. Moreover, TRAP staining for osteoclastogenesis assay confirmed that Col2+ cells cannot differentiate into osteoclasts.

### Col2 negative cells from calvaria bone shows strong unipotent osteogenic potential

According to the lineage tracing results in Figure 3C, both Col2+ cells and Col2− cells are present in calvarial bone. To further test whether Col2− cells also have the differentiation potential, POB (primary osteoblasts) were isolated from the calvaria of Col2-cre; tdTomato (control) and Col2-cre; DTA^fl/−^; tdTomato newborns. We found that approximately 50% of cells were Col2+ in the control group, but few Col2+ cells were detected in the Col2-cre; DTA^fl/−^; tdTomato group, confirming the effectiveness of the Co2+ cell ablation (Figure 7A). Moreover, we found that the cells from control calvarial bone can differentiate into tri-lineage including osteoblasts, chondrocytes and adipocytes. Very interestingly, Col2− cells cannot differentiate into chondrocytes and adipocytes, but have much strong potential to differentiate into osteoblasts compared to the control cells (Figure 7B). Additionally, the wound healing assay showed that Col2− cells have stronger cell migration ability than the control cells (Figure 7C, D).

**Figure 7.**
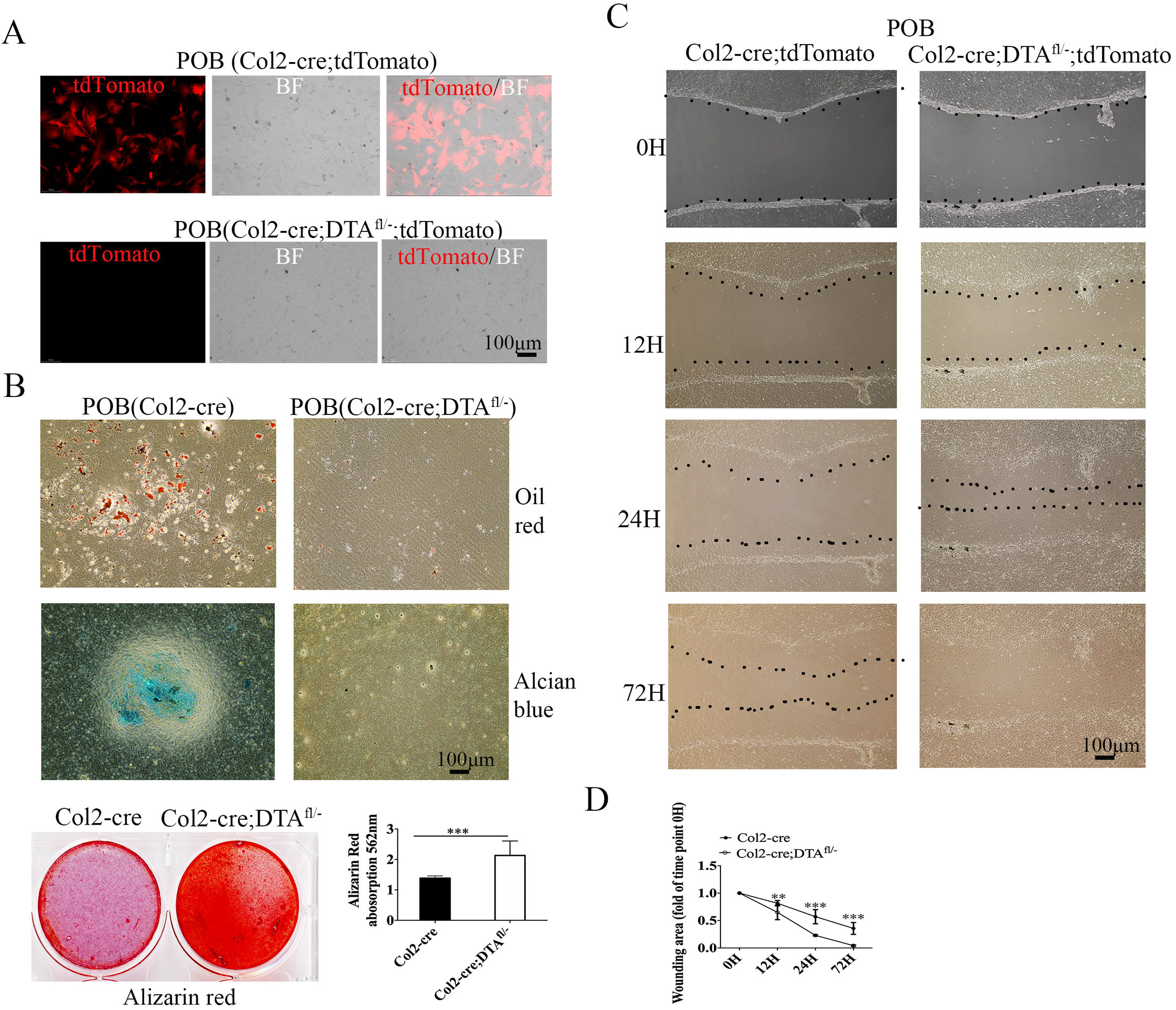
Col2 negative cells from calvaria bone show strong unipotent osteogenic potential. (A) Merged pictures showing the Col2+ POBs cultured from calvaria of Col2-cre;tdTomato and Col2-cre;DTA^fl/−^;tdTomato newborn. Note that: few Col2+ cells can be detected in Col2+ cell ablation group which confirming the effectiveness of the cell ablation technique. (B) Trilineage differentiation assay of cells cultured from Col2-cre and Col2-cre;DTA^fl/−^ new born calvaria bone. (n=3 mice per condition from 3 independent experiments). (C) The different migration ability of POBs cultured from Col2-cre and Col2-cre;DTA^fl/−^ newborn. (D) Quantitative measurements of migration ability in (C). (n=3 mice per condition from 3 independent experiments). All data are reported as the mean ± s.d. Statistical significance was determined by one-way ANOVA and Student’s t-test. *P < .05, **P < .01, ***P < .0001. NS = not statistically significant.

### Col2+ progenitors contribute to chondrocyte differentiation and blood vessel formation during fracture healing

To test whether Col2+ cells contribute to chondrocyte differentiation and blood vessel formation at postnatal stages, we created a closed femoral fracture with intramedullary nail fixation model in 10-week-old mice as described^19^ to observe fracture healing. We first induced tdTomato expression in Col2+ cells in Col2-creERT; tdTomato mice with TM injection at P3 and then subjected the mice to fracture at 10 weeks of age before harvesting tissues after 2 weeks for fracture healing. In the non-injured mice, the contralateral nonfractured femur exhibited prominent tdTomato expression in the AC and metaphysis but little signal in bone marrow (Figure 8A). However, strong tdTomato expression was detected throughout the fracture callus, including both bony and cartilaginous regions. Immunofluorescent staining for CD31 and Col2a1 showed that 95.5% CD31+ and 43% Col2a1+ cells overlapped with Col2+ cells in the callus area of Col2-creERT; tdTomato mice (Figure 8B, C). Additionally, X-ray result showed that ablation of Col2+ cells blocks fracture healing in Col2-creERT; DTA^fl/fl^ mice compared to Col2-creERT (Supplementary Figure7). These results indicate that Col2+ cells play an important role during fracture healing.

**Figure 8.**
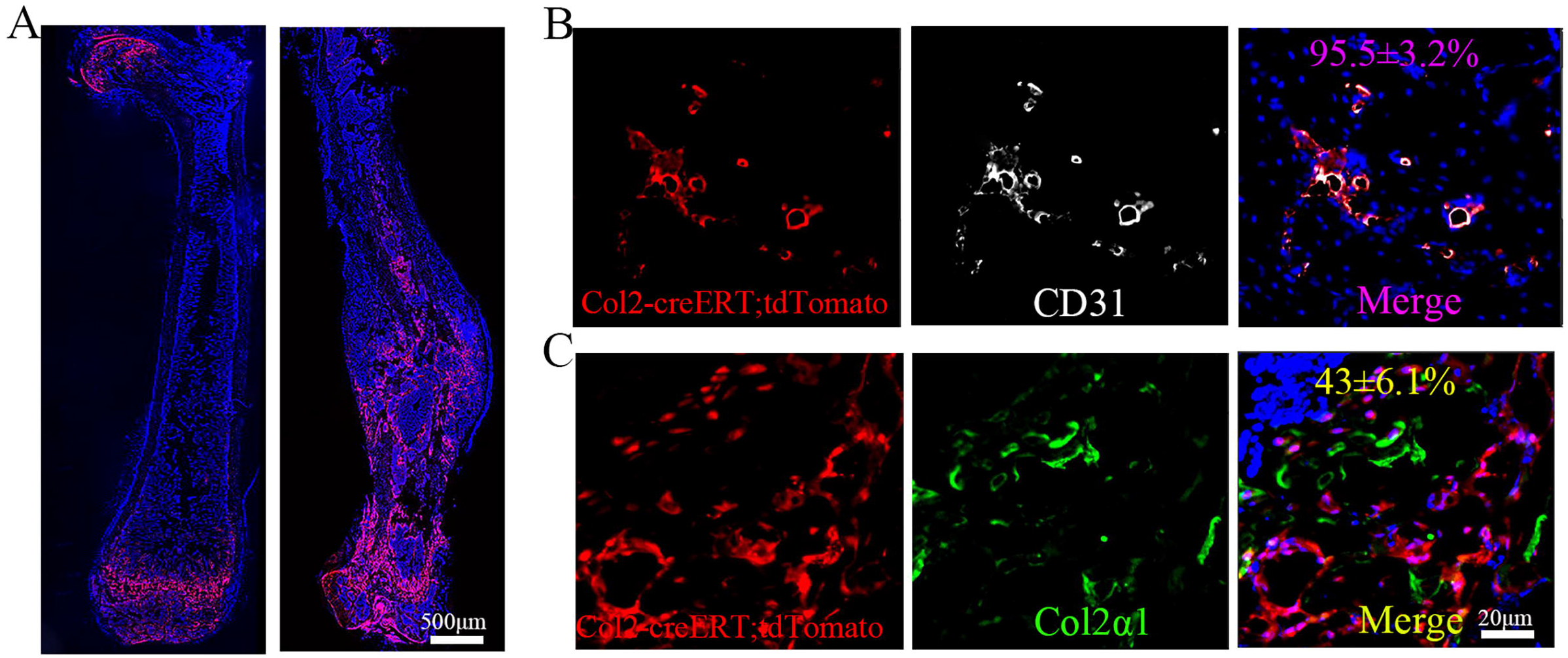
Col2+ progenitors contribute to chondrocyte differentiation and blood vessel formation during fracture healing. (A) The illustration of the experiment design and representative images of femoral section of control and fractured Col2-creERT; tdTomato mice. (It was performed by injecting 75mg/kg tamoxifen into P3 age and then subjected the mice to fracture at 10 weeks of age, before harvesting them after 2 weeks of healing) (B) Representative confocal image of femoral sections through fracture site, with immunofluorescence staining of CD31 (white) to demarcate blood vessel. (n=3 mice per condition from 3 independent experiments). (C) Immunofluorescence staining of Col2a1 (green) on bony callus to demarcate chondrocyte. (n=3 mice per condition from 3 independent experiments). All data are reported as the mean ± s.d. Statistical significance was determined by one-way ANOVA and Student’s t-test. *P < .05, **P < .01, ***P < .0001. NS = not statistically significant.

## Discussion

Previous study from Ono et al ^11^made significant contribution demonstrating that Col2+ cells provides early mesenchymal progenitors to contribute to multiple mesenchymal lineages and continued to provide descendants for regulating the bone growth. This study for the first time further revealed that ablation of Col2+ embryonic progenitors cause mouse lethality at newborn due to dyspnea and absence of skeleton. Additionally, this study provided the evidence that Col2+ cells contributes other important organs’ development including spleen, kidney, heart, et al. Moreover, different from previous concept, we found that both intramembranous and endochondral ossification contribute to long bone and flat bone development. Through genetic lineage tracing, we found that Col2+ cells have multilineage differentiation potential including osteoblasts, chondrocytes, adipocytes and CD31+ blood vessel endothelial cells. Col2+ cells significantly participate in blood vessel formation in several organs including calvaria bone, brain, heart and skin. Postnatal Col2+ progenitors are not only located in bone marrow and GP but also in AC. Additionally, the numbers and differentiation ability of those Col2+ progenitors decrease during aging.

It remains an ongoing debate as to whether bone ossifies endochondrally or intramembranously in mice^20–23^. In craniofacial bones, it has been reported that bones of the cranial base and caudal cranial vault ossify endochondrally, while facial skeleton and rostral cranial vault ossify intramembranously^14,22,23^. Consistently, we found that intramembranous ossification exists in most craniofacial bones and in part of the clavicle. However, different from previous reports^14,24,25^, we found that both intramembranous and endochondral ossification can occur during craniofacial bone development, including the mandible, premaxilla, parietal, frontal, jugal, palatine and temporal bones and clavicle. For example, clavicle was categorized to intramembranous bone which ossify directly from preosteogenic condensations of multipotent mesenchymal cells. However, we found that ablation of Col2+ cells also impaired development of both lateral and medial end of the clavicles, suggesting that both intramembranous and endochondral ossification occur during clavicle development. Besides, the ribs were previously reported to ossify endochondrally^14^. However, we found that ablation of Col2+ cells did not completely disrupt rib formation with a small piece of ribs existed suggesting that ribs ossified both intramembranously and endochondrally. It is well known that long bones, such as the femur and the tibia, ossify endochondrally during embryonic limb development. The existence of bone consisting of Col2− cells in the hindlimbs of Col2-cre; DTA^fl/−^ mice suggested that long bone ossifies both intramembranously and endochondrally during development. Thus, this study provides the first evidence that intramembranous ossification occurs during long bone development, although its function in long bone development needs to be further investigated in the future. The vertebral bone and long bone were previously thought to undergo similar endochondral ossification processes^15,26^. However, the complete loss of the spine in Col2-cre; DTA^fl/−^ mice suggests that vertebral bone ossifies purely endochondrally and Col2+ cells are major stem cells to drive vertebral development.

Through the genetic lineage tracing studies, we found that embryonic Col2+ cells can contribute to most cells in AC, GP, trabecular or cortical bone, tendons, and ligaments as well as some cells in the brain, eye, heart, testicle, epididymis, lung, livers, spleen and kidney. Consistently, ablation of Col2+ cells at embryonic stage in Col2-cre; DTA^fl/−^ mice caused that the mice die at newborn with pale skin, dyspnea, the absence of skeleton and decreased blood vessels, and the severe developmental defects in the heart, kidney^27^. Previous fundamental study has shown that Col2+ cells provide early mesenchymal progenitors in growing bones^10,11^. Consistently, we also found that Col2+ cells have potential to generate many cell lineages, including chondrocytes, osteoblasts, stromal cells and adipocytes. Furthermore, this study provides new evidence that Col2+ cells also contribute fetal organ development including heart, kidneys and blood vessels.

SSCs is a type of somatic stem cell which dedicates to bones. SSCs are considered to play important roles in the development, homeostasis, and regeneration of bone tissues^28^. They are generally defined as self-renewing cells with the “trilineage” potential to differentiate into chondrocytes, osteoblasts and adipocytes. To date, there are three types of SSCs depending on their location: bone marrow-derived SSCs, GP SSCs (a cartilaginous structure separating the primary and secondary ossification centers in growing bones) and perichondrial/periosteal SSCs (the connective tissue surrounding bone)^3,18^. Previous studies have revealed that Col2+ cells in bone marrow, GP and perichondrial/periosteal behave as SSCs^3,29^. Here, we reported, for the first time, that AC also houses a class of Col2+ progenitors. Interestingly, we found that Col2+ cells in AC are similar to Pthrp+ cells ^30^ and Col2+ cells^3,31^ in GP, and few can differentiate into adipocytes *in vitro*. Most interestingly, Col2+ in GP has a stronger osteogenic and chondrogenic differentiation ability than that in articular chondrocytes and BMSCs. The application of Col2+ GP cells, instead of BMSCs, to treat bone defects could be a promising new strategy in the future.

It was reported that the numbers of pthrpr1+ columns in the GP gradually decreased with aging^30^. Consistently to this study, we found that Col2+ cells in articular chondrocytes decreased in age dependent manner^32^. Specifically, the decrease of Col2+ progenitors does not occur at a constant rate with aging; rather, Col2+ progenitors decrease rapidly during 2-3 weeks of age and more slowly at later stages. The drastic lessening of Col2+ cells between 2-3 weeks of age suggests that genetic changes or the activation/deactivation of hormones/growth factors occurs during this stage. Comparing the gene expression changes of cells between 2 and 3 weeks to earlier stage could provide informative mechanisms contributing to bone and cartilage regeneration and repair, and age-related osteoarthritis.

Blood vessels are made up of several different cell types. The inner layer of blood vessels is composed of endothelial cells (ECs), which are covered on the outer, abluminal surface by perivascular (or mural) cells^33^. Angiogenesis requires extensive coordination between the different vascular cell types to ensure that new vessels are fully functional and stable. Increasing evidence indicates that blood vessels form and become specialized in an organ-specific fashion, controlled by local microenvironmental signals and leading to specific molecular signatures in ECs^34^. By lineage tracing of Sox10+ cells, Wang et al^34^ first found that Sox10+ cells only contribute to small numbers of blood vessels in the lung, spleen and kidney and have no contribution to liver vasculature during normal development. Consistently, our result showed that Col2+ cells contribute to CD31+ blood vessel differently in different organs or tissues. Notably, all Col2+ cells were CD31+ in blood vessels of the brain, eyeball, heart, skin and skull bone (Figure 8D, Supplementary figure 5A-D). However, approximately 25.4% of CD31+ were Col2+ cells in the long bone, while no CD31+ cells were Col2+ in kidney vasculature during development.

Bone development during endochondral and intramembranous ossification is tightly coupled with blood vessel branching and growth (angiogenesis), indicating that variations in angiogenesis may be a plausible basis for variations in osteogenesis. However, the cellular resources of blood vessels during bone development remain unclear^26,33,35^. Previously studies suggested that blood vessels can form via two processes. In early embryogenesis, mesodermal cells differentiate into hemangioblasts (the progenitors of ECs and blood cells), which migrate to specific locations and aggregate to form the first primitive vessels in a process termed vasculogenesis^36^. Subsequently, new blood vessels arise by the process of angiogenesis-the expansion of existing vascular networks through a series of processes such as EC sprouting, migration, proliferation, vessel anastomosis and pruning ^37^. Our findings revealed, for the first time, that blood vessels also originate from Col2+ progenitors. Moreover, our *invitro* osteoclasto-genesis assay by culturing the sorting Col2+ cell and Col2− cells from bone marrow further confirm Col2+ progenitor cannot differentiate to osteoclasts, suggesting that the blood vessel forming cells and blood cells in bone marrow originated from different cell lineage.

The critical role of angiogenesis during endochondral and intramembranous ossification has been well recognized^14^. Although similarities exist between calvarial and long bone osteogenesis in vascular invasion into surrounding avascular loose mesenchyme and the association of invading vessels with mineralization, fundamental difference in the osteogenesis of endochondral and intramembranous bones hints the hypothetical differences in angiogenesis in the two types of bone formation^33^. In agreement with this theory, our result showed that almost all of Col2+ cells contributes to calvaria bone, however, parts of Col2+ cells contribute to long bone blood vessel, suggesting the difference in cell origins in the development of blood vessels during the two types of bone development. Interestingly, although our results suggest lower numbers of Col2+ blood vessels in long bone during normal development than those in skull bone, the number of Col2+ blood vessels increases upon cell expansion following fracture, which is consistent with a previous report on Sox10+ cells^34^.

Osteoblasts are the cells responsible for new bone formation during skeletal development, remodeling, and repair. To date, much progress has been made in defining the molecular and cellular properties of the osteoblast phenotype through characterization of primary calvarial-derived osteoblast cultures from chicks, rats, and mice^24,38^. We report that Col2+ cells were present in calvarial-derived cells when we isolated POBs from mouse calvaria according to a previously described protocol^39^. Interestingly, calvarial-derived cells from Col2-cre control mice can differentiate into osteoblasts, chondrocytes, and adipocytes. However, Col2− cells from Col2-cre; DTA^fl/−^ mouse calvaria can only differentiate into osteoblasts. This result suggested that Col2+ cells in the calvaria might be progenitor cells with multipotent differentiation ability. Although great efforts have been made to identify the difference between intramembranous and endochondral ossification, intramembranous osteoblasts have not been successfully isolated from bone^15^. In our study, by comparing Col2− intramembranous osteoblasts from Col2-cre;DTA^fl/−^ and Col2+ endochondral osteoblasts from Col2-cre mice, we found that intramembranous osteoblasts have stronger osteogenic ability than endochondral osteoblasts. Further experiments are required to establish the relationship between Col2− and Col2+ osteoblasts in the future.

In summary, our study, for the first time, revealed that both endochondral and intramembranous ossification are involved in most calvarial flat bone and long bone development. Col2+ cells are major progenitors to contribute to skeletal development and CD31+ blood vessel endothelial development in many organs, including calvariae bone, long bone development and fracture repair. Number and differentiation ability of Col2+ progenitors decrease during aging in mice. Identifying new Col2+ and/or Col2− progenitors from GP, AC and calvaria bone provide new promising therapeutic strategies for bone regeneration and repair as well as treatment of bone and cartilage diseases.

## Method and materials

### Mice

All procedures regarding mouse housing, breeding, and collection of animal tissues were performed as per approved protocols by the Institutional Animal Care and Use Committee (IACUC) of the University of Pennsylvania, in accordance with the IACUC’s relevant guidelines and regulations. All animals were of the C57BL strain, and all mice were housed in specific pathogen-free conditions. Col2-cre^40^, Col2-creERT^41^, R26-tdTomato^42^, and DTA^fl/fl^ ^43^ mice were obtained from Jackson Laboratory (Bar Habor, ME, USA). Col2-cre; DTA^fl/−^ and Col2-creERT; DTA^fl/fl^ mice were generated by breeding DTA^fl/fl^ mice with Col2-cre or Col2-creERT mice. Col2-creERT; DTA^fl/fl^ were injected with TM at the indicated time points to induce the postnatal deletion of Col2+ cells. To monitor the topography of Col2-cre and Col2-creERT function, we crossed Col2-creERT and Col2-cre mice with R26-tdTomato reporter mice and analyzed the tdTomato labeled patterns in long bone, cartilage, and knee joint at different time points. Mice were euthanized by overdosage of carbon dioxide. TM (T5648, Sigma) solution preparation and administration were performed as previously described^44^. Briefly, TM was first dissolved in 100% ethanol (100 mg/mL) and then diluted with sterile corn oil to a final concentration of 10 mg/ml. The TM-oil mixture was stored at 4 °C until use. Mice in both the control and experimental groups were administered the same dose of TM (75 mg/kg body weight) according to the time points defined in the experimental protocol.

### Histology

Samples were dissected under a stereo microscope to remove soft tissues, fixed in 4% paraformaldehyde for overnight at 4 °C, and then decalcified in 10% ethylenediaminetetraacetic acid (EDTA) for 14 days at 4 °C. Decalcified samples were cryoprotected in 30% sucrose/PBS solutions and then in 30% sucrose/PBS:OCT (1:1) solutions overnight at 4 °C. Samples were then embedded in an OCT compound (4583, Sakura) under a stereo microscope and transferred to a sheet of dry ice to solidify the compound. Embedded samples were cryosectioned at 6μm using a cryostat (CM1850, Leica).

### Safranin O/fast green staining

Mouse tibia were sectioned and stained with Safranin O/fast green staining to visualize cartilage and assess proteoglycan content, as described previously^45^. Samples on slides were stained with Weigert’s iron hematoxylin and fast green and then stained with 0.1% Safranin O solution.

### Alizarin red/Alcian blue staining

Alizarin red/Alcian blue staining was used to stain the whole skeleton as reported previously^44^. Briefly, the skeletons of newborn mice (n=3) were fixed with 90% ethanol and then stained with 0.01% Alcian blue solution (26385-01, Electron Microscopy Sciences) and 1% Alizarin red S solution (A47503, Thomas Scientific). Stained skeletons were stored in glycerol.

### Von Kossa staining

Von Kossa staining was performed with 1% silver nitrate solution (LC227501, Fisher Scientific) in a glass coplin jar placed under ultraviolet light for 20-30 minutes^46^. Unreacted silver was washed with 5% sodium thiosulfate (01525, Chem-Impex). The slides were dehydrated with graded alcohols and mounted with permanent mounting medium.

### Immunofluorescence microscopy

Tibia sections with a thickness of 6 μm were gently rinsed with PBS and incubated with proteinase K (20 μg/mL, D3001-2-5, Zymo Research) for 10 min at room temperature. Subsequently, sections were blocked in 5% normal serum (10000 C, Thermo Fisher Scientific) in PBS-T (0.4% Triton X-100 in PBS) or incubated with antibodies against type II collagen (1:100, ab34712, Abcam) and CD31 antibody (1:100, sc-81158, Santa Cruz Biotechnology) in blocking buffer at 4 °C overnight. Tissue sections were washed 3 times with PBS and then incubated respectively with the secondary antibody of Alexa Fluor 488-conjugated anti-rabbit (1:200, A11008, Invitrogen) and Alexa Fluor 647-conjugated anti-mouse (1:200, A-21236, Invitrogen) antibodies for 1 hr at room temperature. Coverslips were mounted with Fluoroshield (F6057, Sigma-Aldrich).

To quantify the percentage of tdTomato+ cells, multiple fields of Z-stacked pictures were randomly captured. At least 30 images were measured. The percentage of tdTomato+ cells was calculated from the ratio of tdTomato+ cells to total cells observed in each compartment and each sample (five sections were collected for each sample). Six mice were evaluated in each group. Assessments were independently performed by two authors who were blinded to the groups. The average percentage of tdTomato+ cells in each sample was pooled and calculated by two authors. The average tdTomato+ percentage in each group from six mice was pooled and calculated.

### PEGASOS passive immersion and blood vessel staining

PEGASOS passive immersion and blood vessel staining approaches were performed to stain the blood vessel as reported previously.^47^ Briefly, before transcardiac perfusion, mice were anesthetized with an intraperitoneal injection of a combination of xylazine and ketamine anesthetics (xylazine 10-12.5 mg/kg; ketamine, 80-100 mg/kg body weight). Then, 50 ml ice-cold heparin PBS (10 U/ml heparin sodium in 0.01 M PBS) was injected transcardially to wash out the blood. Finally, 50 ml 4% paraformaldehyde (PFA) in 0.01 M PBS (pH 7.4) was infused transcardially for fixation.

For clearing the calvarial bones, samples were fixed in 4% PFA at room temperature for 12 h, and then, were immersed in 0.5M EDTA (pH 7.0) at 37 °C in a shaker for 2 days. Next, samples were decolorized with the Quadrol decolorization solution for 1 day at 37 °C in a shaker. Then transfer samples into 1.5ml Eppendorf tubes containing blocking solution composed of 10% dimethyl sulfoxide, 0.5% IgePal630 and 1X casein buffer in 1ml 0.01M PBS for blocking overnight at room temperature. After blocking, stain samples with CD-31 primary antibodies in blocking solution for three days at 4°C on a shaker. Wash samples at least three times with PBS at room temperature for one day. Put samples into freshly prepared secondary antibodies diluted with blocking solution for another three days at 4°C on a shaker. Wash the samples again for half a day. Samples were then placed in 30%, 50%, 70% gradient tB delipidation solutions for 4 hours each and then tB-PEG for 2 days for dehydration. Samples were immersed in BB-PEG medium at 37 °C for half a day for clearing. Images were acquired by Sp8 confocal microscope (Leica) with the lens of 25X. 3-D reconstruction images were generated using Imaris 9.0 (Bitplane). Stack images were generated using the “volume rendering” function. Optical slices were obtained using the “orthoslicer” function. 3-D images were generated using the “snapshot” function.

### Micro-CT analysis

Quantitative analysis of the gross bone morphology and microarchitecture was performed by Micro-CT (Micro-CT 35, Scanco Medical AG, Brüttisellen, Switzerland) at Penn Center for Musculoskeletal Disorders (PCMD), University of Pennsylvania. Briefly, the femurs from 4-week-old Col2-creERT and Col2-creERT; DTA^fl/fl^ mice were fixed, scanned and reconstituted as three-dimensional images. The cross-sectional scans were analyzed to evaluate the changes in the femur. The ROIs were then compiled into 3D data sets using a Gaussian filter (sigma=1.2, support=2) to reduce noise and were converted to binary images with a fixed grayscale threshold of 316. The trabecular bone architecture was assessed between the distal femoral metaphysis and midshaft. Two hundred (200) slices (2 mm) above the highest point of the growth plate were contoured for trabecular bone analysis of the bone volume fraction (BV·TV^−1^), trabecular thickness (Tb. Th), trabecular number (Tb.N), trabecular separation (Tb. Sp).

### Cell culture

Primary osteoblasts (POBs) will isolated from newborn cavaria bone. Briefly, calvaria bones were dissected from newborns, subjected to sequential collagenase type II (2 mg/ml) (17101015, Gibco) and trypsin (SH3004202, GE Healthcare) digestion for 30 min and cut into tiny pieces. The minced bones were treated with collagenase type II and trypsin again before being plated in tissue culture dishes. POBs that migrated from the bone pieces were passaged and used for the further studies.

Bone marrow stem cells (BMSCs) were isolated as previously described^39^. Briefly, femurs and tibias were dissected from 4-week old Col2-cre; tdTomato mice. The bones were cut on both ends with sterile blades (22079690, Fisher Scientific). The bone marrow was flushed with complete α-MEM using 23-gauge needle syringes. The cells were then transferred for flow cytometry sorting and cell culture.

GP chondrocytes was isolated as previously described^30^. Briefly, distal epiphyses of femurs from 4-week old Col2-cre; tdTomato were manually dislodged and attached soft tissues and woven bones were carefully removed. Dissected epiphyses were incubated in 3 ml Hank’s balanced salt solution (HBSS, H6648, Sigma) at 37 °C for 60 min on a shaking incubator. AC and secondary ossification centers were subsequently removed. Dissected GPs were minced using a disposable scalpel (22079690, Fisher Scientific) and further incubated with Liberase TM at 37 °C for 60 min on a shaking incubator. Cells were mechanically triturated using an 18-gauge needle and a 1 ml Luer-Lok syringe and filtered through a 70 μm cell strainer into a 50 ml tube on ice to obtain a single-cell suspension. Cells were pelleted and resuspended in appropriate medium for subsequent purposes.

AC chondrocytes was isolated as previously described^19^. Briefly, 4-week old Col2-cre; tdTomato mice were euthanized at postnatal day 28 (P28). The AC from the femoral heads was isolated by first thoroughly removing the soft tissue and bone, followed by incubation with collagenase type IV (LS004188, Worthington Biochemical Corp) (3 mg/mL) for 45 min at 37 °C. Cartilage pieces were obtained and incubated in 0.5 mg/mL collagenase type IV solutions overnight at 37 °C. Cells were then filtered through a 40 μm cell strainer, collected and obtained a single-cell suspension. Cells were pelleted and resuspended in appropriate medium for subsequent purposes.

### Flow cytometry

Cells were resuspended in 100 ml of cell staining buffer (420201, Biolegend), and incubated with anti-CD16/32 antibody for 20 minutes on ice for blocking Fc receptors, followed by staining with fluorochrome-conjugated or isotype control antibodies on ice for 20 minutes. To identify MSCs, the cells were incubated with anti-CD31-APC (102419, Biolegend, 1:100) antibody, and then stained with biotin-conjugated antibodies. After washing with staining medium, the cells were incubated with streptavidin-brilliant violet 421 TM (Biolegend, 1:500). Flow cytometry analysis was performed using a five-laser BD LSR Fortessa (Ex. 405/488/561/640 nm) and FACSDiva software. Acquired raw data were further analyzed using FlowJo software (Tree Star). Representative plots of at least three independent biological samples are shown in the figure.

### CFU-F assay and *in vitro* differentiation

The following procedures were modified from a previous report^10^. For CFU-F assays, freshly prepared unfractionated bone marrow single-cell suspensions were plated at a density of ~10^4^ cells/cm ^2^ (0.21 ml of culture medium) in 100-mm dishes in DMEM supplemented with 20% FBS qualified, 10 M Y-27632 (TOCRIS) and 1% penicillin/streptomycin. For CFU-F assays with sorted cells, tdTomato+ cells were sorted and directly seeded into culture at a density of 10 cells/cm^2^ in 6-well plates, ensuring that colonies would form at clonal density and therefore could be counted. The cultures were incubated at 37 °C in a humidified atmosphere with 5% O_2_ and 10% CO_2_ for 7-10 days. CFU-F colonies were counted after 7-10 days of culture.

For the *in vitro* differentiation assay, tdTomato+ cells were sorted and seeded into each well of 48-well plates and cultured for 14 days. Adipocyte, chondrocyte and osteoblastic differentiation was induced with different differentiation media and detected by staining with Oil red (O0625-25G, Sigma-Aldrich), Alcian blue (26385-01, Electron Microscopy Sciences) and Alizarin red (A47503, Thomas Scientific), respectively as performed previously^48,49^.

### Bone fracture

Closed femoral fractures with intramedullary nail fixation were created in 10-week-old mice (Col2-creERT;tdTomato or Col2-creERT;DTA^fl/fl^;tdTomato) as described^19^. Briefly, closed fractures were generated by a three-point blunt guillotine driven by a dropped weight, which creates a uniform transverse fracture of the femur. The fractured femurs were harvested at 12-week-old for analysis. n=6 mice per condition from 3 independent experiments.

### Scratch-wound assay

Scratch-wound assay did as previously described^50^. The same numbers of POBs cells from Col2-cre;tdTomato and Col2-cre; DTA^fl/fl^;tdTomato mouse calvaria bone were seeded in six-well plates. The monolayer cells were scratched and then cultivated under normal conditions. The migration distances at 0, 12, 24, and 72 h were measured after scratching for each group.

### Statistics

All data are presented as the mean±s.d. The Shapiro-Wilk test for normality and Bartlett’s test for variance were performed to determine the appropriate statistical tests. Student’s t-test for the comparison between two groups or one-way ANOVA followed by Tukey’s multiple comparison test for grouped samples was performed. The number of animals and repetitions of experiments are presented in the figure legends. The program GraphPad Prism (GraphPad Software, Inc., San Diego, USA) was used for these analyses. (*P<0.05, **P<0.01, ***P<0.0001. NS=not statistically significant.)

## ACKNOWLEDGMENTS

Research reported in this publication was supported by the National Institute of Dental and Craniofacial Research and the National Institute of Arthritis and Musculoskeletal and Skin Diseases, and National Institute on Aging, part of the National Institutes of Health, under Award Numbers DE023105, AR066101 and AG048388 to S.Y. XL was supported by China Scholarship Council (CSC) Grant #201706260178. The content is solely the responsibility of the authors and does not necessarily represent the official views of the National Institutes of Health.

## AUTHOR CONTRIBUTIONS

XL, DJ and SY performed the experiments, interpreted the data and wrote the initial draft of the manuscript. STY managed mice colonies and assisted with the experiments. SY conceived, supervised the study and wrote the manuscript. LQ and HZ provided critical suggestions, reagents and technical assistance during the study.

