## Supplemental figures for "Type II collagen-positive progenitors are major stem cells to control skeleton development and vascular formation"

The authors declare no conflict of interest.

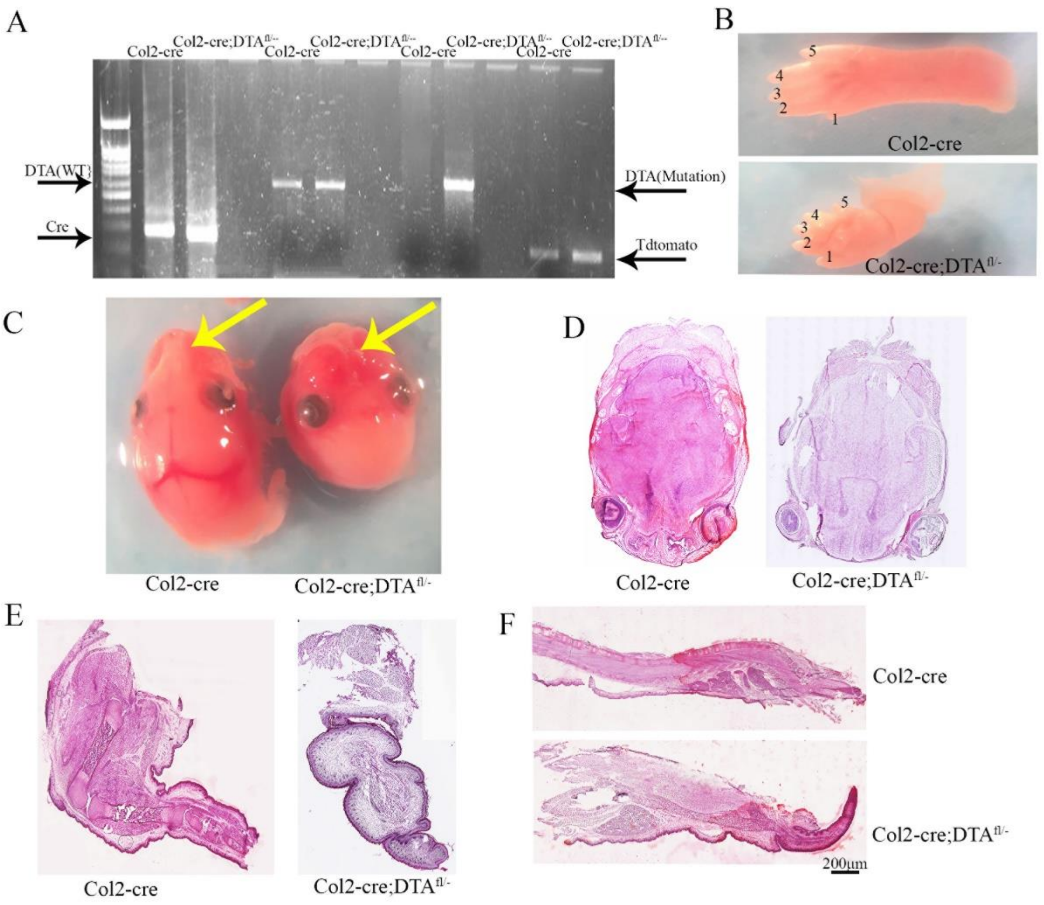


**Supplementary Figure 1. Genotyping and histology analysis of skeleton in newborn**

(A) The detection of different band in genotyping for Col2-cre;tdTomato and Col2-cre;$DTA^{fl/-}$;tdTomato mice. (n=3 mice per condition from 3 independent experiments). (B) Gross appearance the toes of Col2-cre and Col2-cre;$DTA^{fl/-}$ newborn. (C) Gross appearance the skull in Col2-cre;tdtomato and Col2-cre;$DTA^{fl/-}$ embryos at P0. Yellow arrow: the cleft palate in mutant mice. (D) H&E staining of coloron section in the skull of Col2-cre and Col2-cre;$DTA^{fl/-}$ mice. (E) H&E staining the low extremities in Col2-cre and Col2-cre;$DTA^{fl/-}$ mice. (F) H&E staining of the sagittal section of spine in Col2-cre and Col2-cre;$DTA^{fl/-}$ mice. n=3 mice per condition from 3 independent experiments.

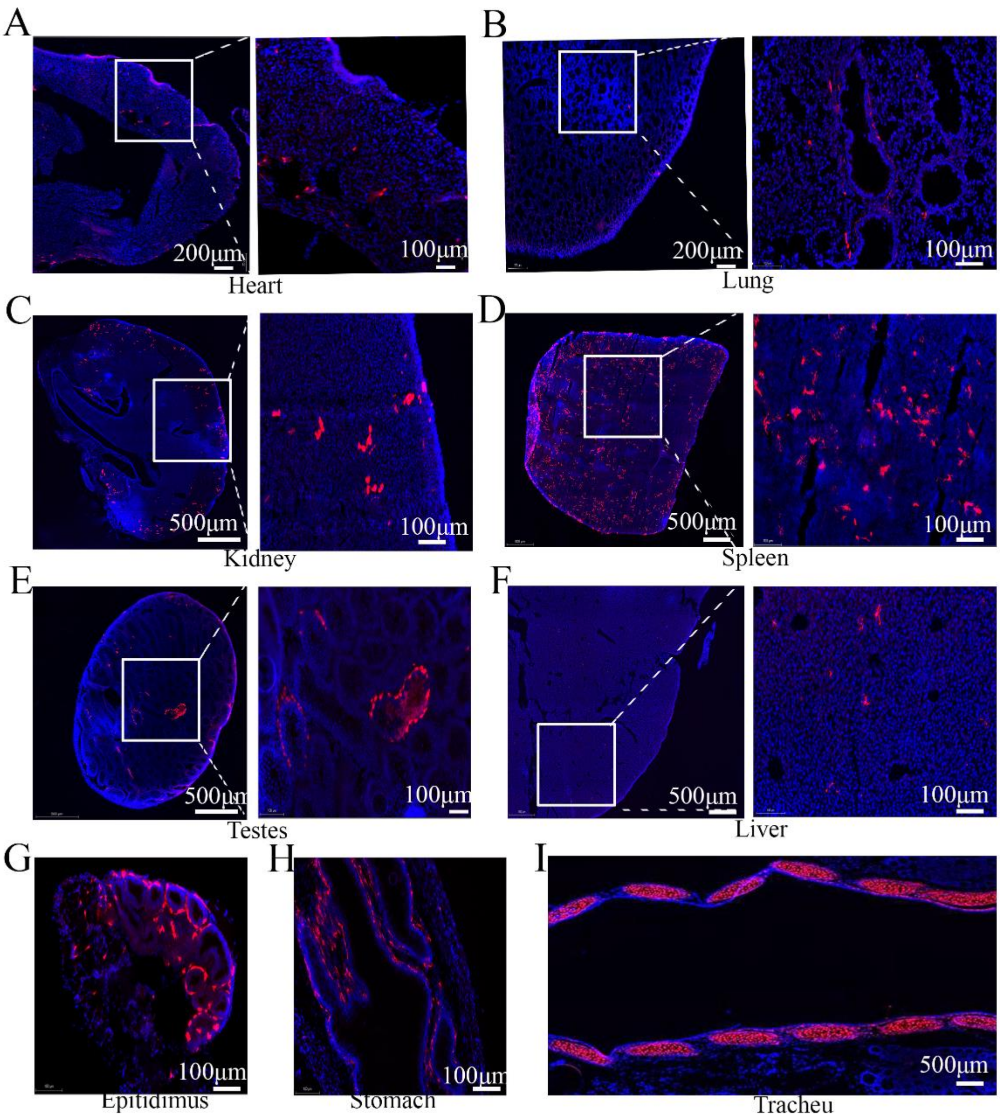


**Supplementary Figure 2. Col2+ cells contribute to many organs' development**

Representative images from lineage tracing of Col2-cre+ in heat (A), Lung (B), Kidney (C), Spleen (D), Testes (E), Liver (F), Epitidimus (G), Stomach (H), Tracheu (I) in 4-week-old Col2-cre; tdTomato mice. n=3 mice per condition from 3 independent experiments.

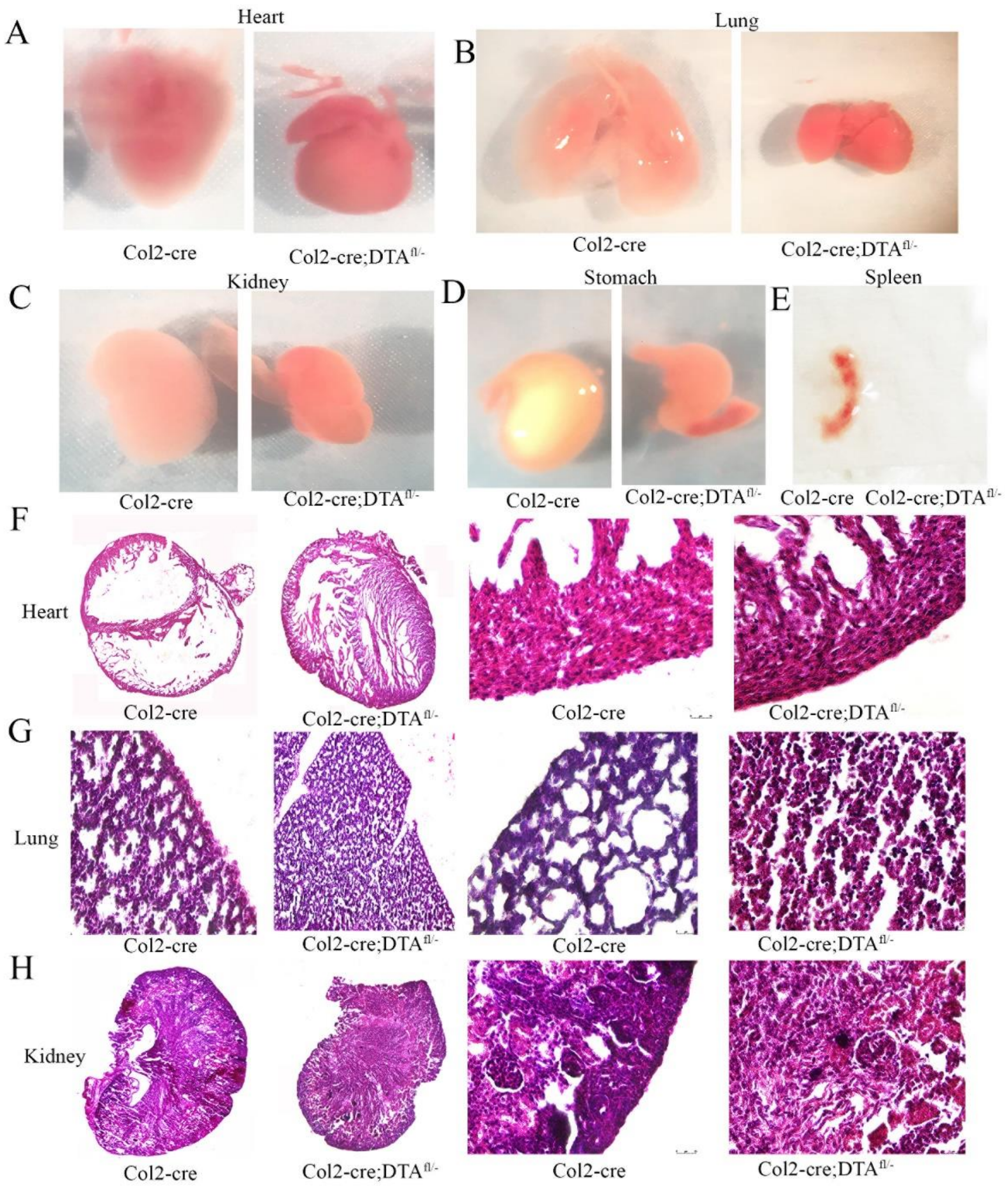


**Supplementary Figure 3. Histology analysis different organs in newborn**

Gross appearance the Heart (A), Lung (B), Kidney (C), Stomach (D), Spleen (E) of Col2-cre and Col2-cre; DTA$^{fl/-}$ mice at P0. Note that the spleen completely loss in mutant mice. H&E staining analysis of the heart (F), Lung (G), Kidney (H) of Col2-cre and Col2-cre; DTA$^{fl/-}$ mice at P0. n=3 mice per condition from 3 independent experiments.

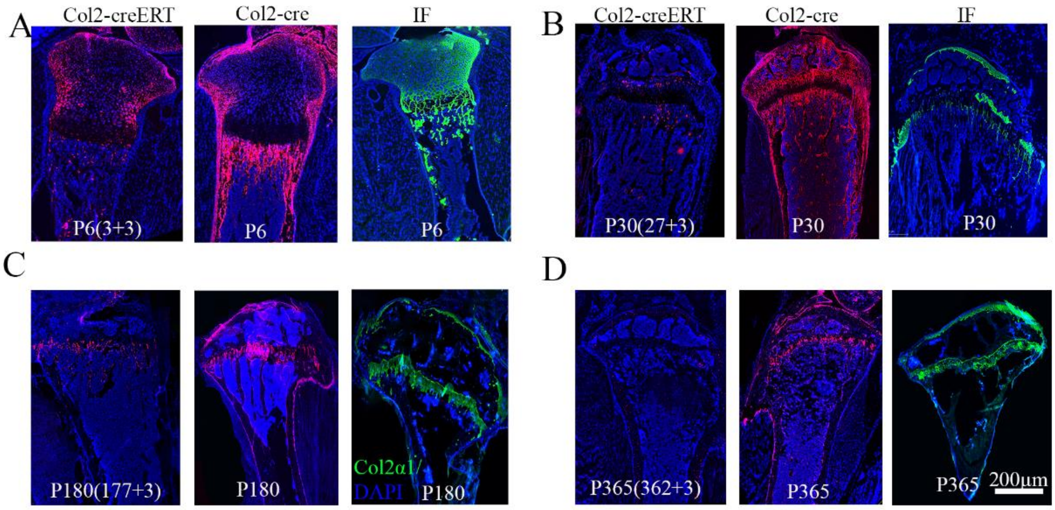


**Supplementary Figure 4. The inconsistence of Col2+ cells and type II collagen expression in long bone**

The distribution of Col2+ cells by lineage tracing for Col2-creERT;tdTomato and Col2-cre;tdTomato mice and the collagen 2 expression (Col2a1) detected by IF staining in long bone at P6 (A), P30 (B), P180 (C), P365 (D). n=3 mice per condition from 3 independent experiments.

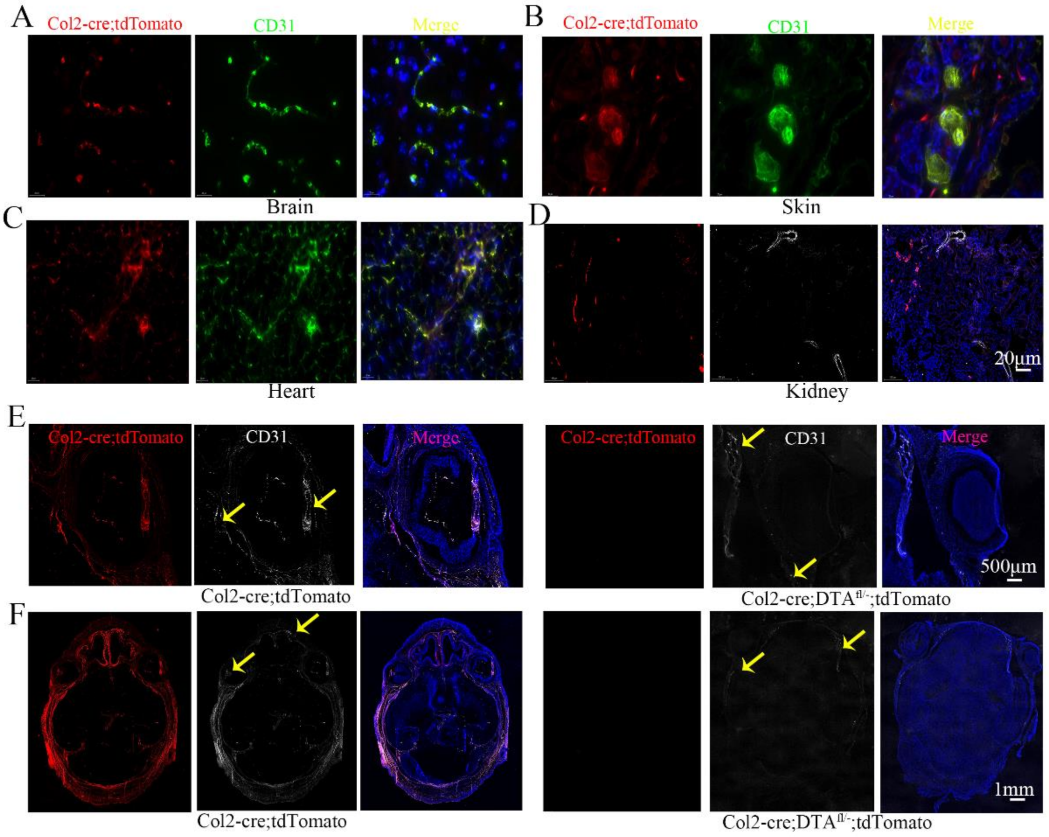


**Supplementary Figure 5. The distinct contributions of Col2+ cells to CD31+ vascular development in different organs**

(A) Representative immunofluorescence staining of CD31 (green) on brain tissues to demarcate blood vessel. (B) Representative immunofluorescence staining of CD31 (green) on skin tissues to demarcate blood vessel. (C) Representative immunofluorescence staining of CD31 (green) on heart tissues to demarcate blood vessel. (D) Representative immunofluorescence staining of CD31 (white) on kidney to demarcate blood vessel. (E) Representative immunofluorescence staining of CD31 (white) on eye in Col2-cre; tdTomato and Col2-cre;DTA$^{fl/-}$; tdTomato mice to demarcate blood vessel change. (F) Representative immunofluorescence staining of CD31 (white) on coronal section of skull in Col2-cre; tdTomato and Col2-cre;DTA$^{fl/-}$; tdTomato mice to demarcate blood vessel change. Yellow arrow: the positive CD31 staining. n=3 mice per condition from 3 independent experiments.

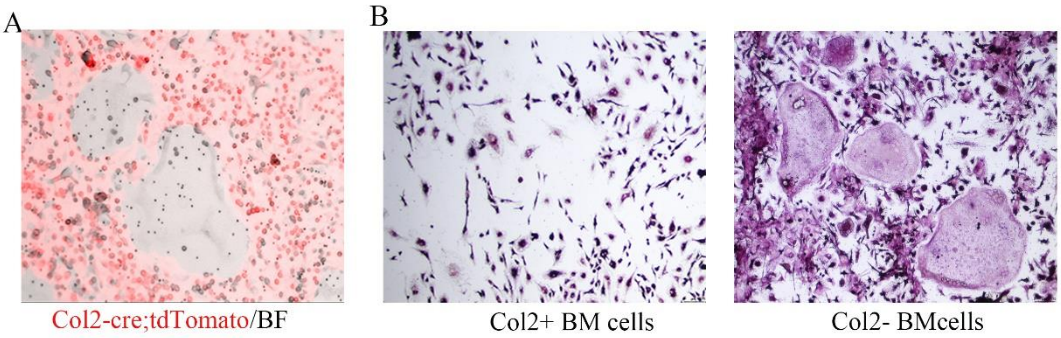


**Supplementary Figure 6. Col2+ cells cannot differentiate to osteoclast**

(A) The overlap bright filed and tdTomato fluorescence in osteoclast from bone marrow in 4-week-old Col2-cre;tdTomato mouse. (B)TRAP staining for osteoclastogenesis assay from Col2- and Col2+ bone marrow cells confirmed that Col2+ cells cannot differentiate into osteoclasts. n=3 mice per condition from 3 independent experiments.

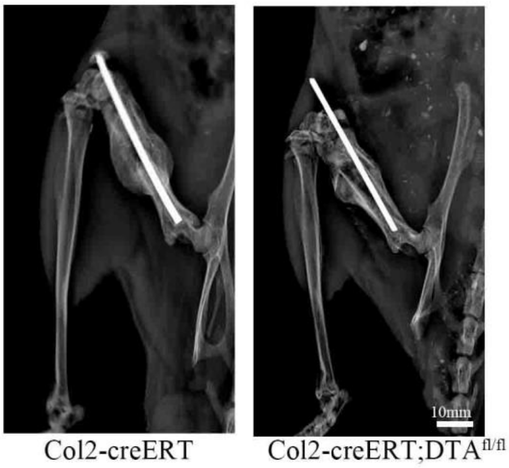


**Supplementary Figure 7. Postnatal deletion Col2+ cells delayed fracture healing**

Representative X ray picture showed the fracture site in 2 weeks following fracture of Col2-creERT and Col2-creERT;DTA$^{fl/fl}$ mouse. n=3 mice per condition from 3 independent experiments.
